# Generation of Rapid Sequences by Motor Cortex

**DOI:** 10.1101/2020.06.09.143040

**Authors:** Andrew J. Zimnik, Mark M. Churchland

## Abstract

Rapid execution of motor sequences is believed to depend upon the fusing of movement elements into cohesive units that are executed holistically. We sought to determine the contribution of motor cortex activity to this ability. Two monkeys performed highly practiced two-reach sequences, interleaved with matched reaches performed alone or separated by a delay. We partitioned neural population activity into components pertaining to preparation, initiation, and execution. The hypothesis that movement elements fuse makes specific predictions regarding all three forms of activity. We observed none of these predicted effects. Instead, two-reach sequences involved the same set of neural events as individual reaches, but with a remarkable temporal compression: preparation for the second reach occurred as the first was in flight. Thus, at the level of motor cortex, skillfully executing a rapid sequence depends not on fusing elements, but on the ability to perform two computations at the same time.

## Introduction

Decades of research have documented the events within motor cortex that accompany execution of a voluntary movement. But what is ‘a movement’? The answer is unambiguous for a single, discrete behavior, such as pressing a button or reaching to a target. The subject begins a trial at rest, moves, then returns to rest. ‘A movement’ is the action that occurs between periods of quiescence. But how does this conception extend to other behaviors? For example, how many movements are used to enter a four-digit bank code? If button presses are separated by considerable time, it seems reasonable to assume they are generated individually. Conversely, when the sequence is well-learned and rapidly executed, it seems likely to be generated as a unified whole.

There is indeed considerable evidence that the temporal separation between sequence elements relates to how that sequence is internally produced^1–3^. When practiced in a specific order, the time between elements becomes minimal, and they are said to form a functional unit: a ‘motor chunk’^1,3,4^. Motor chunk formation occurs spontaneously during long sequences, yielding subsets of swiftly executed elements with longer separations between subsets^1,5,6^. A specific chunk structure can be encouraged by manipulating the delay between elements^7–9^. Thus, chunking is both encouraged by, and a result of, the need to rapidly execute multiple elements in sequence.

The neural and computational basis of this behavioral phenomenon is less clear. Is chunking a ‘cognitive’ skill, relating to how groups of actions are recalled and conveyed to downstream motor areas? Or is chunking a motor skill, in which motor areas generate, holistically, actions that were originally generated separately? Studies of motor cortex report ‘sequence selectivity’: responses that reflect whether an action is performed alone or within a sequence^10–15^ (although see^16–18^). Sequence selectivity concurs with the hypothesis that chunking is a motor phenomenon, with chunks executed differently from their component elements.

Yet sequence selectivity may arise for other reasons. Muscle activity can differ depending on whether an action is performed within a sequence. This can be controlled for, but not perfectly. Sequence selectivity could also result from overlap of execution-related and preparation-related activity^19,20^. If the next element is prepared as the first is being completed, a ‘new’ pattern of activity, not observed during execution of any single element, will be produced. For these reasons, one wishes to compare neural activity not to a null hypothesis (identical responses regardless of whether an element occurs within a sequence) but to predictions made by competing hypotheses. One hypothesis is that rapid production of a sequence does not depend on anything ‘new’ at the level of motor cortex, but simply reflects the sequential preparation and execution of each element. The competing hypothesis is that elements are prepared and executed as a unified whole.

The maturing characterization of neural events during single reaches makes it possible to derive concrete predictions from these hypotheses. A reach involves three distinguishable neural processes observable within motor and premotor cortex. The first is preparatory. Preparatory activity reflects the identity of the pending movement and is proposed to seed subsequent movement-generating neural dynamics^21–26^. The second is a putative ‘trigger signal’: a large change in neural state ~150 ms before movement onset^27,28^. This change does not reflect movement identity but rather the timing of movement initiation. The third is execution-related: richly time-varying activity that arises ~10-20 ms before muscle activity begins^21,22^. If two elements are prepared and executed sequentially, then preparation-, triggering-, and execution-related events should occur twice. If two elements are prepared and executed holistically, each event should occur once. These competing hypotheses also make different predictions regarding the structure of neural activity during these stages. For example, before the sequence begins, preparatory activity should reflect only the first element under the sequential hypothesis, but should reflect aspects of both elements under the holistic hypothesis.

We recorded from premotor and primary motor cortex in monkeys trained to execute sequences of two reaches, either in rapid succession (compound reach) or separated by an imposed pause (delayed double-reach). As expected, the imposed pause produced clearly sequential preparation and execution: the prepare-trigger-execute motif was observed twice. Unexpectedly, sequential preparation and execution persisted during compound reaches. This was true even though the full sequence was known in advance and had been practiced tens of thousands of times. It was true despite a very rapid pace, in which muscle activity for the second reach began as the first was landing. Neural responses revealed how this rapid pace was possible. Preparation for the second reach occurred during execution of the first reach and did so without disrupting the first reach. Thus, at the level of motor and premotor cortex, skilled performance depends not on fusing elements, but upon the ability to prepare one element while executing another.

## Results

### Task and Behavior

We trained two rhesus macaques (monkeys B and H) to perform a modified delayed-reach task (Fig. 1a,b). All trials began with a randomized (0-1000 ms) instructed delay period. Each trial required the monkey to make either a single reach, two reaches separated by an instructed pause (delayed double-reach), or two reaches with no pause between them (compound reach). Target color indicated which should be acquired first. A salient visual cue (the diameter of the colored portion of the first target) indicated whether a pause was required. For monkey H, the pause between delayed double-reaches was always 600 ms. For monkey B it was variable (100, 300, or 600 ms; the last is used for most analyses) and indicated by the visual cue. Thus, all key information – target locations and any instructed pause – was given during the instructed delay. Reach paths are illustrated (Fig. 1a) for all single reaches, for three compound reaches that began down-and-right, and for three compound reaches that began down-and-left (Extended Data Fig. 1 shows paths for all compound reaches).

**Fig. 1.**
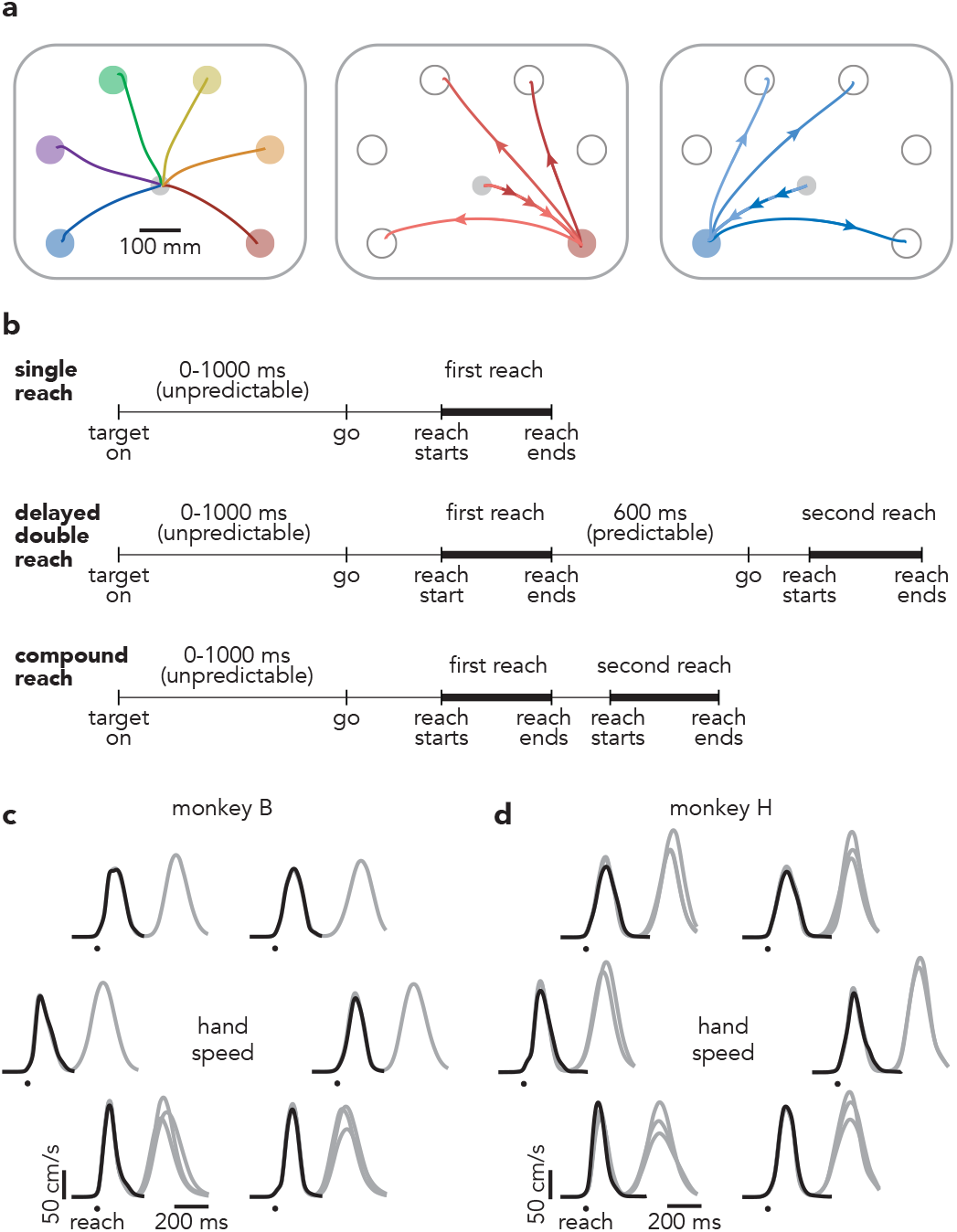
Task structure. **a,** Mean hand paths for all single reach conditions and a subset of compound reach conditions. *Left panel*: single reaches. *Middle panel*: three compound reaches that began with a reach to the bottom-right target. *Right panel*: three compound reaches that began with a reach to the bottom-left target. **b,** Trial timing. Single reaches employed a variable instructed delay period. Delayed double-reaches employed an instructed pause between reaches, the duration of which was indicated during the initial instructed delay. Compound reaches had no instructed pause and monkeys reached for the second target immediately after the first was touched. **c,** Mean hand speed for all single reach (*black*) and compound reach (*grey*) conditions. Traces are grouped according to the location of the first target. Every target location served as the first target for at least one compound reach condition; some were used for three. Averages were calculated across all trials and recording sessions. Data are for monkey B. **d,** Same as **c** but for Monkey H. Every target location served as the first target for two or three compound reach conditions.

Compound reaches were performed briskly; the hand stayed on the first target only briefly before moving to the second target. Median dwell times on the first target were 119 ms (monkey B) and 137 ms (monkey H). The median duration for the full two-reach sequence was 561 ms (monkey B) and 645 ms (monkey H). This rapid pace resulted from extensive training over months, with each sequence performed tens of thousands of times. This pace is even faster than that reported in other motor sequence tasks performed by non-human primates, where dwell times are typically >200 ms^10,11,29^.

To enable comparisons at the neural level, every compound reach condition shared the same first target with a matched single-reach condition. Comparisons at the neural level are most interpretable if the reaches themselves (not just target locations) are well-matched. This was indeed the case. Reach-speed profiles (Fig. 1c,d) were very similar for single reaches (black) and the first reach of compound reaches (gray); correlations were 0.99 ± 0.002 (mean ± standard deviation, monkey B) and 0.98 ± 0.03 (monkey H). The most noticeable difference was that the first reach of a compound reach tended to be slightly faster than the corresponding single reach: by 3 ± 2.8% and 3 ± 9.4% (monkey B and H, change in peak velocity averaged across conditions). We also examined the activity of the major muscles of the upper arm and shoulder (Fig. 2a and Extended Data 3 show two such recordings). Muscle activity began changing ~100 ms before reach onset (*circles*). During the subsequent 275 ms, muscle activity during compound reaches was similar to that during single reaches to the same target (*gray* and *black* traces overlap). Mean correlations were 0.93 ± 0.15 and 0.93 ± 0.15 (monkey B and H, average across muscles and conditions). Comparing compound reaches with matched single reaches, muscle activity magnitude was, on average, slightly lower for monkey B (2 ± 15%) and slightly higher for monkey H (8 ± 22%). Thus, behavior approached the desired ideal: an identical first reach regardless of a second reach. The match was even closer when comparing delayed double-reaches with single reaches, both for velocity (ρ = 0.99 ± 0.0004 and 0.99 ± 0.001 monkey B and H) and muscle activity (average ρ = 0.98 ± 0.05 and 0.99 ± 0.05).

**Fig. 2.**
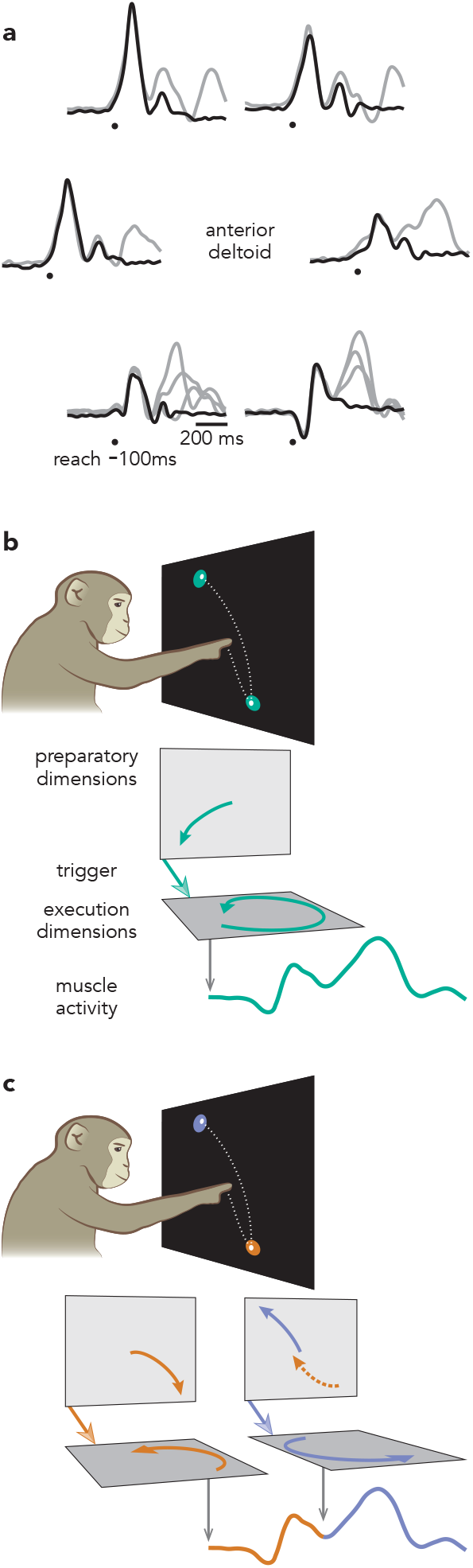
Potential strategies for generating compound reaches. **a,** Mean EMG for single (*black*) and compound (*grey*) reach conditions. Data are from the deltoid of Monkey B. Traces are grouped according to the location of the first target. **b,** Illustration of the holistic strategy. Under the holistic strategy, compound reaches are produced as single movements, which require only a single pattern of preparatory activity and a single trigger signal. **c,** Illustration of the sequential strategy. Under the sequential strategy, compound reaches are generated by preparation, triggering, and execution occurring twice in succession.

The final relevant comparison is between compound and delayed double-reaches, which would ideally have identical second reaches. This is probably impossible; during compound reaches the second reach is generated while muscle activity is still in flux after the first reach (e.g., post-reach co-contraction). Consistent with this, second-reach muscle activity was strongly but imperfectly correlated between compound and delayed double-reaches: ρ = 0.90 ± 0.16 and 0.83 ± 0.25 (monkey B and H, average across muscles and conditions).

During compound reaches, generation of the second reach requires muscle activity, at some point, to depart strongly from that during single reaches. This occurred ~275 ms after initial EMG onset. This translates to ~25 ms before the end of the first reach and ~150 ms before the onset of the second reach. This timing underscores how rapidly compound reaches unfolded – muscle activity began changing in anticipation of the second reach as the first reach was ending. As a result, muscle activity during compound reaches evolved continuously throughout the full sequence; there was no pause or plateau between reaches.

### Basic Properties of Neural Responses

We recorded activity from the arm region of motor cortex (dorsal premotor cortex and surface primary motor cortex, Extended Data Fig. 2) using 32-channel linear-array electrodes. Recordings consisted of well-isolated single neurons and high-quality multi-unit isolations (227 and 587 units from monkey B and H). As in prior studies^10–13,30^ many units exhibited sequence selectivity – i.e., responses associated with a reach could depend on whether it was part of a sequence. Comparing delay-period activity before matched single versus compound reaches, 11% and 24% of recorded units (monkey B and H) showed significantly different responses during at least one pair of conditions (p < 0.001, unpaired t-test, adjusted for multiple comparisons). These percentages grew when comparing movement-epoch activity: to 38% and 60% (when considering time-averaged activity) or 79% and 95% (when considering the full temporal pattern; see *Methods*).

Such differences are potentially consistent with a holistic strategy, which requires that an individual reach be generated ‘differently’ when part of a sequence. Had we found no significant differences, a holistic strategy could therefore have been rejected. Yet the mere presence of significant differences does not discriminate between sequential and holistic strategies. Muscle activity also differs slightly, rendering small but potentially significant neural differences unavoidable. Furthermore, sequence selectivity is potentially expected during movement even under the sequential hypothesis. If preparation for the second reach overlaps with execution of the first, that would create a ‘new’ pattern of activity not observed during any single reach. Thus, distinguishing between holistic and sequential strategies requires testing more specific predictions.

### Derivation of Predictions

Such predictions are possible because of extensive prior characterization of neural activity in motor cortex during single reaches^19,20,31–34^. This has yielded a paradigm in which reach generation is proposed to be subserved by three internal processes: preparation, triggering, and execution^21–26,35–37^. Preparatory activity, typically observed during an instructed delay, has long been hypothesized to be a necessary precursor of voluntary movement^19,20,31^. Importantly, preparatory activity also occurs without an instructed delay^21,31,35,38^ and can develop very rapidly^21,35^. Preparatory activity is proposed to seed execution-related activity and thus specify the identity of the executed reach. The transition from preparatory to execution-related activity coincides with a large condition-invariant change in the neural state, proposed to result from a movement-triggering input^27,39–41^. Traditionally it has been challenging to discriminate preparation-related activity from execution-related activity when they occur close in time^31^. However, the recent focus of neural subspaces^42–44^ – sets of neural dimensions that capture prominent response features – opens up new approaches. In particular, preparation-related, triggering-related, and execution-related activity occur in nearly orthogonal subspaces^21,27,28,36^ and can thus be separated via linear projections of the population response.

The above paradigm makes it possible to concretely specify the competing hypotheses and derive their predictions. Under the holistic strategy (Fig. 2b) the two elements of a compound reach are produced cohesively, as if they were a single movement. An appropriate preparatory state is established, a trigger signal arrives, and execution-related dynamics generate activity that produces the desired continuous pattern of muscle activity. Under the sequential strategy (Fig. 2c), a compound reach is simply two distinct movements, prepared and executed sequentially. The first is prepared, triggered, and executed as if it were a single reach. Shortly thereafter, these three stages occur a second time. Continuous muscle activity results from the concatenated output.

The holistic and sequential hypotheses yield three mutually exclusive predictions. First, preparatory activity should occur once if the holistic strategy is employed and twice if the sequential strategy is employed. Second, under the holistic strategy, preparatory activity before a compound reach should differ from that before a corresponding single reach. In contrast, the sequential strategy predicts that preparatory activity will be similar, because only the first reach is prepared at that time. Third, the holistic strategy predicts a single trigger-related change in activity, whereas the sequential strategy predicts that each reach will be preceded by its own trigger-related event.

While it is hoped that observations will consistently obey all the predictions of one hypothesis, it is also possible for results to be mixed or fail to follow any of the above predictions. Thus, evaluating these predictions provides not only a test of the two hypotheses, but of the paradigm itself. Partly for this reason, our task also included delayed double-reaches as a reference. The imposed pause between the two reaches, instructed during the initial delay, is expected to dissuade the use of a holistic strategy. Delayed double-reaches thus afford an opportunity to confirm the predictions of the sequential strategy in a context where it is likely to be used. This provides a foundation for asking what occurs when there is no pause and the holistic strategy becomes viable.

### Timing of Single-Neuron Preparatory Activity

Most of the above predictions must be tested at the population level because most neurons exhibit a mixture of preparation-related, triggering-related, and execution-related activity. Yet a small percentage of recorded neurons displayed nearly ‘pure’ preparatory activity, affording an opportunity to tentatively explore some predictions. Consider the response illustrated in Figure 3a. For single-reach conditions, delay-period activity is strongly selective: high before reaches to a down-and-left target and low before reaches to a down-and-right target. Selectivity collapses by movement onset. This pattern held across all conditions (Extended Data Fig. 4) but for simplicity is shown here for the two target locations that were also used for the first reach of delayed double-reaches. (For monkey B, delayed double-reaches used fewer combinations than compound reaches, to allow different inter-reach pause durations while maintaining reasonable trial-counts.) The response of this example neuron is consistent with participation in preparatory computations but not execution-related computations. This interpretation is supported by responses during delayed double-reaches (Fig. 3b). Due to the imposition of a 600 ms pause between reaches, one expects monkeys to adopt a sequential strategy: the first reach should be prepared during the initial instructed delay, and the second should be prepared during the imposed pause. Consistent with that expectation, this neuron is selectively active during the initial instructed delay, largely silent during the first movement, and active again during the imposed pause (Fig. 3b). The second bout of (presumed) preparatory activity has different selectivity from the first, as expected given that the second reach differed from the first. In both cases, the pattern of putatively preparatory activity collapses as muscle activity starts changing (traces at bottom).

**Fig. 3.**
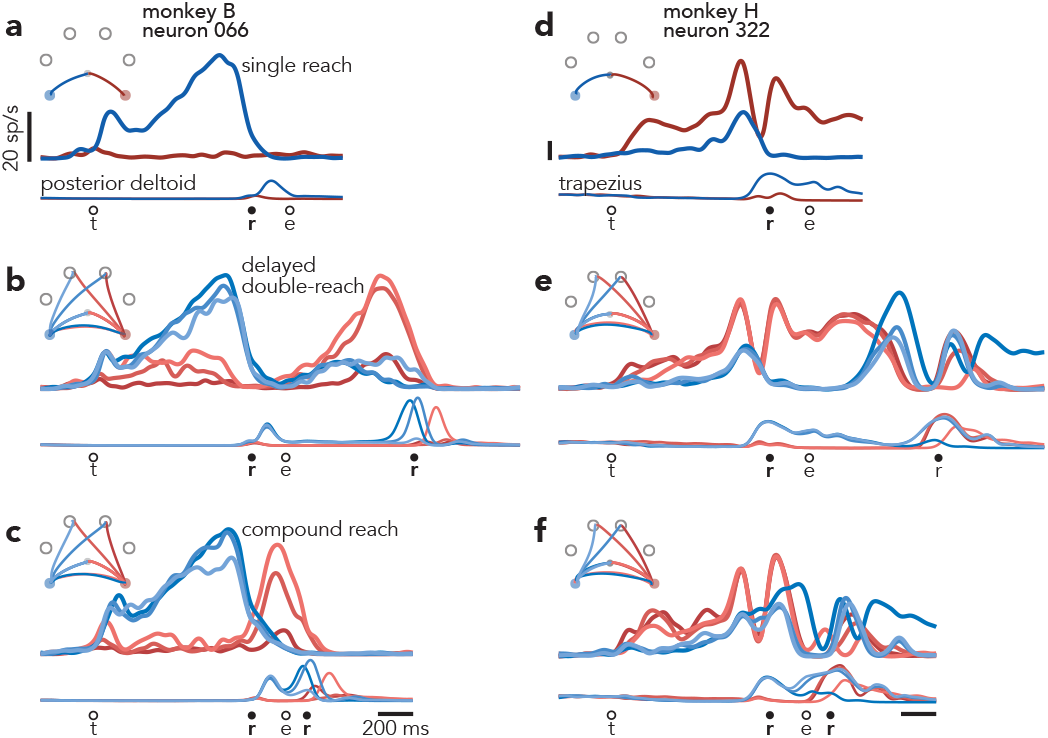
Responses of single neurons. **a,** Activity of one neuron, recorded from monkey B during single reaches to bottom-right (*red*) and bottom-left (*blue*) targets. Thin traces at bottom plot the activity of the posterior deltoid. Circles indicate the time of target onset (*t*), reach onset (*r*), and reach end (*e*). All traces are trial-averages of data that have been aligned twice: once to target onset and once to reach onset. See *Methods* for a complete description of data pre-processing. **b,** Response of the same neuron and muscle during delayed double-reach conditions. Monkey B performed six such conditions, all of which are shown. **c,** Response of the same neuron and muscle during compound reaches. Trace color corresponds to the direction of the first reach. Monkey B performed ten compound reach conditions, but for illustration only those with corresponding delayed double-reach conditions are shown. Data for all conditions is plotted in Extended Data Fig. 4. **d,** Activity of one neuron, recorded from monkey H during single reaches. Same format as **a**. Thin traces at bottom plot the activity of the trapezius. **e,** Activity of the same neuron and muscle during delayed double-reaches (same format **b**). **f,** Activity of the same neuron and muscle during compound reaches. Responses during all tested conditions are plotted in Extended Data Fig. 5a.

Compound reaches were executed so rapidly that the two epochs of muscle activity fused (Fig. 3c, *bottom traces*). However, the neuron’s response did not shift to the pattern predicted by the holistic strategy. That hypothesis predicts that preparatory activity during the instructed delay should reflect the full two-reach movement, and thus differ from the preparatory activity before single reaches. This was not observed. The three compound reaches that began with a down-and-left reach (Fig. 3c, *blue traces*) shared similar firing rates during the instructed delay, despite differing second reaches. Indeed, these firing rates were similar to that when preparing only a down-and-left reach (Fig. 3a). Analogously, activity was low for the three compound reaches that began with a down-and-right reach (*red traces*), in agreement with activity for a single down-and-right-reach. Furthermore, the neuron was active not only before the first reach, but again before the onset of the second reach, suggesting a second bout of preparation. More broadly, this neuron’s response pattern during compound reaches (Fig. 3c) was similar to that during delayed double-reaches (Fig. 3b), but with a much earlier second peak. During compound movements, the second peak occurred during the first movement rather than during the pause between movements. This neuron was recorded from monkey B, for whom delayed double-reaches were also examined with imposed pauses of 300 ms and 100 ms (in addition to the 600 ms pause in Fig. 3b). Inspection of this neuron’s response across pause durations (Extended Data Fig. 4) confirms that the compound-reach response lies on a continuum with the delayed-double reach responses. Thus, the responses of this example neuron suggest that not only is the sequential strategy employed during delayed double-reaches (as expected), it may continue to be employed during compound reaches.

Of course, little can be concluded from the response of one neuron. Not only might that response not be representative, it allows examination of only a subset of the key predictions. A challenge when considering all neurons is that most are active during both preparation and movement. Such responses are typically complex even during single reaches (Fig. 3d) and more so during delayed double-reaches (Fig. 3e) and compound reaches (Fig. 3f). Indeed, once one considers all conditions (only a subset of which are shown in Fig. 3) the responses of most neurons are bafflingly complex (Extended Data Fig. 5) and it becomes nearly impossible to address simple questions – such as whether compound movements involve two bouts of preparation – via inspection. Testing the competing predictions of the sequential and holistic strategies thus depends upon separating preparatory and execution-related signals at the population level.

### Time-course of Preparatory Activity

Preparatory and execution-related signals were recently shown to be separable at the population-level^21,36^. The correlation structure between neurons is essentially unrelated during preparation versus movement. Thus, the neural dimensions that best capture preparatory activity are nearly orthogonal to those that best capture execution-related activity. Preparatory and execution-related signals can therefore be separated via projection onto appropriate sets of orthogonal dimensions. Identifying those dimensions requires leveraging task epochs where preparation is occurring without execution and vice versa. Our task affords a few such epochs. There is no movement during the instructed delay or the pause between delayed double-reaches. Yet the pending reach is presumably being prepared at these times. Conversely, movement execution is (by definition) occurring during single reaches, each delayed double-reach, and the final reach of compound reaches – times when preparation is unlikely because there is no immediately pending next movement. We used activity from these epochs to define 20 preparatory and 20 execution dimensions. This approach carries a caveat: dimensions optimized to capture preparation before one set of reaches may sub-optimally capture preparation before different reaches. As an abstract example, dimensions optimized to capture preparation for slow reaches are expected to imperfectly capture preparation for fast reaches. This matters because the second compound reach is driven by modestly different patterns of muscle activity (see above) from the corresponding second delayed double-reach. Thus, any preparation for the second compound reach may be incompletely captured by the preparatory dimensions, as they were optimized to capture preparation for modestly different movements. This limitation is acceptable so long as one is conservative when interpreting measures of variance captured.

Projecting population data onto the first preparatory dimension (Fig. 4a-c) yields a response similar to a single-neuron PSTH (with greater symmetry due to mean-centering during preprocessing, see *Methods*). This resemblance is expected: the activity patterns captured by these dimensions are building blocks of single-neuron responses. For example, the first preparatory dimension was the primary contributor (with a large negative weight) to the response of the example neuron in Fig. 3a-c, accounting for 92% of its across-condition response structure. Of course, this purity was rare; most neurons contained sizeable contributions from multiple dimensions, both preparatory and execution-related. This highlights the utility of the projections, which can be relatively pure even when single neurons are not.

**Fig. 4.**
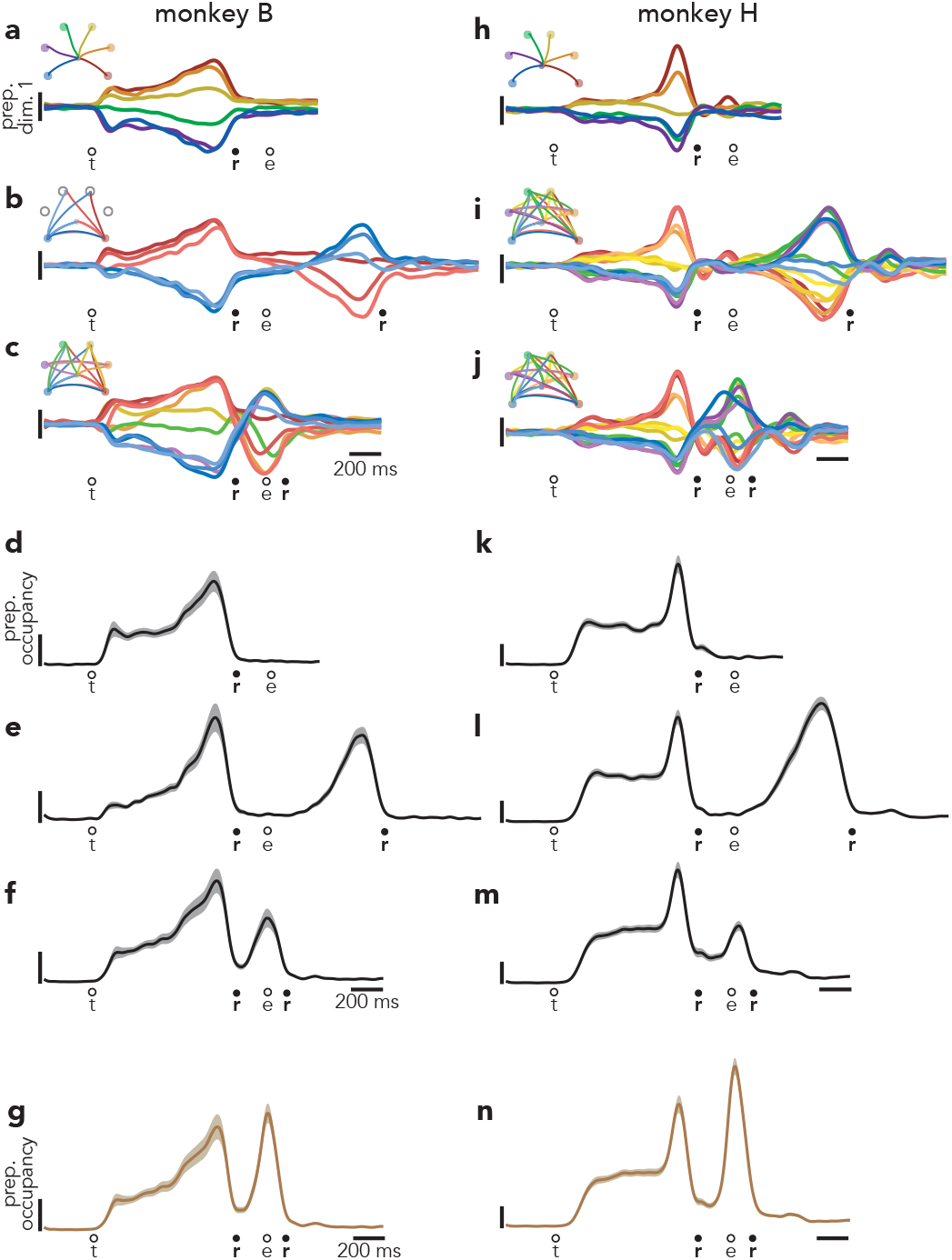
Time-course of activity in preparatory dimensions. **a,** Projection of population activity, during single reaches, onto the first preparatory dimension. Data are from monkey B. Unlike in Fig. 5, data are shown for all conditions. Vertical scaling is arbitrary and conserved across panels. **b,** The same projection for delayed double-reaches. **c,** The same projection for compound reaches. **d,** Preparatory subspace occupancy (for all 20 dimensions) during single reach conditions. Shaded regions indicate the standard deviation of the sampling error (equivalent to the standard error of the mean) estimated by resampling. As occupancy is a measure of normalized firing rates, the units are arbitrary. **e,** Preparatory subspace occupancy during delayed double-reaches. **f,** Preparatory subspace occupancy during compound reaches. **g,** Same as **f**, but preparatory dimensions were found using an expanded range of times that included activity during the brief dwell-period between compound reaches. **h-j**, Same as **a-c** but for monkey H. Note that monkey H performed more two-reach conditions than monkey B. **k-n**, Same as **d-g** but for monkey H. Occupancy just prior to the second reach in compound reach conditions (**f,m**) was significantly greater than the occupancy during the same time period of single reaches for both monkeys (p < 0.001 for both monkeys via a bootstrap procedure; see *Methods*).

During delayed double-reaches, activity in the first preparatory dimension is expected to obey the predictions of the sequential strategy. This was indeed the case (Fig. 4b). There were two bouts of strong selectivity: one before the first reach (peaking just before movement onset) and another before the second reach (again peaking just before movement onset). Activity before the first reach was similar to that before the corresponding single reach. *Red traces* cluster together and resemble the response before a single down-and-right reach. *Blue traces* cluster together and resemble the response before a single down-and-left reach. During the second bout of preparation, the order of the traces largely inverts, in agreement with the physical reversal of the reach (when the first was rightwards, the second had a leftwards component).

During compound reaches, activity in the first preparatory dimension (Fig. 4c) obeyed the predictions of the sequential strategy and violated the predictions of the holistic strategy. There were two bouts of preparatory subspace activity. The first occurred during the instructed delay. The second peaked just as the first movement ended. Furthermore, there was no evidence that the first bout of preparation reflected the two-reach sequence. The ten traces in Fig. 4c are color coded by the identity of the first reach. The pattern of preparatory activity before the first reach was nearly identical to that before single reaches (Fig 4a).

The full pattern of activity during compound reaches was very similar to that during delayed double-reaches (which had four fewer conditions) with the exception of timing. For delayed double-reaches, the second bout of preparation occurred when the hand was stationary during the imposed pause. For compound reaches, the second bout of preparation occurred essentially contiguously with the decline of the first bout. Thus, the second bout of preparatory dimension activity was developing while the hand was still in flight. Because monkey B performed delayed double-reaches with different instructed pauses, we could ask whether activity during compound reaches lay on a continuum with those conditions. In other words, is the pattern of activity similar, but with the second bout of preparation simply occurring earlier? This was indeed the case (Extended Data Fig. 6). Thus, activity in the first preparatory dimension was consistent with the hypothesis that, during compound reaches, only the first reach is prepared during the instructed delay period and the second reach is then prepared as the hand is in flight.

To assess activity across all preparatory dimensions, we computed preparatory subspace occupancy: the across-condition variance of the neural state as a function of time. For single-reach conditions (Fig. 4d) the preparatory subspace was occupied during the instructed delay but not during the reach^21,38^. For delayed double-reaches (Fig. 4e; Extended Data Fig. 6g,h,i), the preparatory subspace was occupied twice, once before each reach. This continued to be true for compound reaches (Fig. 4f), consistent with the results observed above for the first preparatory dimension. For compound reaches, the second peak was smaller than the first. This might seem to suggest that the second bout of preparation is weaker. However, a smaller second peak is expected for technical reasons. As discussed above, dimensions optimized to capture preparation before one set of movements are essentially guaranteed to sub-optimally capture preparation for a different set of movements. To be conservative, activity before the second compound reach was not used when optimizing preparatory dimensions (we wished to determine whether a second bout of preparation naturally emerged). If one is less conservative, and allows optimization to also employ the epoch just before the second compound reach, the two preparatory peaks are of comparable size (Fig. 4g, Extended Data Fig. 7).

Results were similar for monkey H (Fig. 4, *right column*). Monkey H performed a larger set of two-reach combinations, all of which were performed both as delayed double-reaches and compound reaches. To prevent trial-counts from becoming overly high, the instructed pause was always 600 ms for delayed double-reaches. Otherwise the task was unchanged. Results were correspondingly similar. Compound reaches involved two bouts of preparation (Fig. 4j,m,n). During the first bout, selectivity matched that before single reaches and before delayed double-reaches (compare order of colored traces across panels h-j).

### Patterns of Preparatory Activity

The analysis above indicates that delay-period activity before compound reaches is similar to that before corresponding single reaches. To explore further, we chose a time-point near the end of the initial bout of preparation: 120 ms before first-reach onset. In the top two preparatory dimensions, the pattern of neural states resembled the spatial layout of the reach targets (Fig. 5a,b). This is true even though preparatory activity is not a literal representation of target location^23^. For compound reaches (*circles*) preparatory states clustered according to first-reach identity. For example, *blue circles* correspond to three compound reaches that all begin with a down-and-left reach. These cluster near the state before a single down-and-left reach (*blue triangle*), despite involving dissimilar second reaches.

**Fig. 5.**
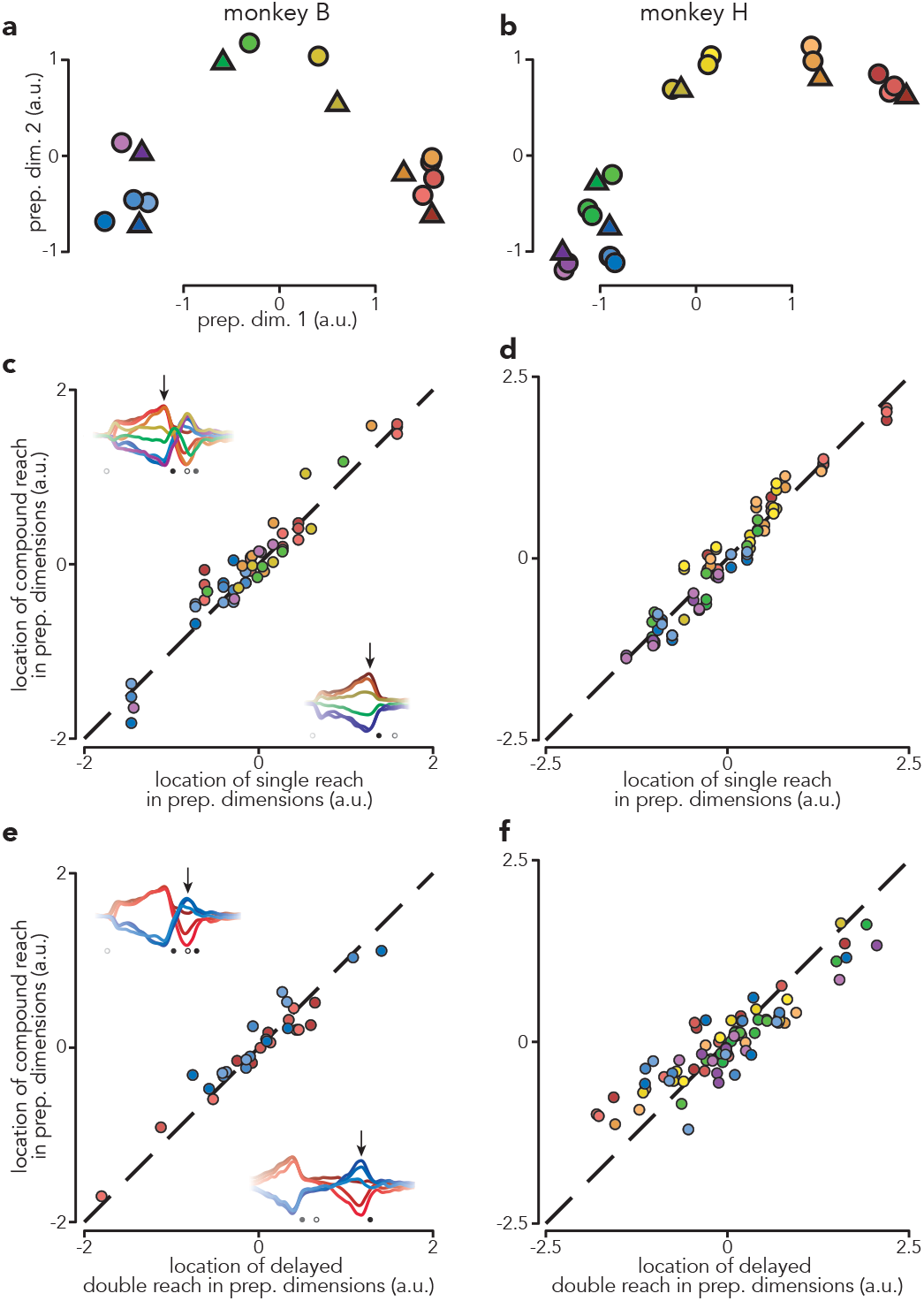
Patterns of preparatory activity. **a,** Projections of population activity, just before first-reach onset, onto the top two preparatory dimensions. Triangles represent single reach conditions and circles represent compound reach conditions. Marker color indicates first-reach direction. Data are from monkey B. **b,** Same as **a** but for monkey H. Monkey H performed more compound reach conditions than monkey B. **c,** Comparison of preparatory activity, before first-reach onset, between compound and single reaches. Each circle plots the location in preparatory subspace of activity before one compound reach versus that for the corresponding single-reach. Arrows on each inset indicate when preparatory subspace activity was assessed. Each compound-reach condition contributes five data-points – one for each of the top five preparatory dimensions. Dashed line indicates unity slope. Data are for monkey B. **d,** same as **c** but for Monkey H. **e,** Comparison of preparatory activity, prior to the second reach, between compound reaches and matched delayed double-reaches. Arrows on each inset indicate when preparatory subspace activity was assessed. **f,** same as **e** but for monkey H.

Across the top five preparatory dimensions and all conditions, the location of the preparatory state before a compound reach was always similar to that before the corresponding single reach (Fig. 5c,d; ρ = 0.96 for both monkeys; p < 0.0001 for both). We focused on the top five dimensions because each of the additional fifteen captured, on its own, little variance. Nevertheless, results were nearly identical when extended to all twenty preparatory dimensions: ρ = 0.92 and 0.92 for both monkeys (p < 0.0001 for both).

According to our working paradigm, similar preparatory states should lead to similar time-evolving patterns of execution-related activity. To test this, we repeated the above analysis for the execution-related dimensions and for multiple times (Extended Data Fig. 8). Execution-related activity was very similar for the first compound reach and the corresponding single reach: ρ = 0.91 and 0.93 (monkey B and H, p < 0.0001 for both). Thus, the first reach of a compound reach involves preparatory and execution-related neural events very similar to those accompanying a matched single reach.

A prediction of the sequential hypothesis is that not only should compound reaches involve two bouts of preparation, but the second bout should reflect the second reach much as if it were performed in isolation. An ideal test of this prediction is not possible because the second reach of a compound reach is not identical to the second reach of a delayed double-reach; the former is generated while muscle activity from the first reach is still dissipating. Yet while not identical, muscle activity patterns are still fairly similar. The sequential hypothesis thus predicts a considerable similarity between the preparatory state preceding the second compound versus delayed double-reach. This was indeed the case: ρ = 0.96 and 0.90 (monkeys B and H, p < 0.0001, both monkeys). Notably, this was true despite the very different circumstances under which preparatory state was measured: while the first compound reach was in flight versus at the end of the imposed pause between delayed double-reaches.

### Trigger-related signals

The final prediction of the holistic and sequential hypotheses concerns trigger-related events. We have proposed that reaches are generated by a movement-specific preparatory neural state interacting with execution-related dynamics. Yet movement-specific signals, while prevalent, are not the largest signals in motor cortex during reaching. Instead, the dominant aspect of the neural response is independent of reach identity. This ‘condition invariant signal’ (CIS) begins ~150 ms before movement onset^27,28,39,40^ and is proposed to reflect a large input that triggers movement execution. Consistent with this, the dominant response component in mouse motor cortex during reaching depends upon thalamic inputs, and silencing those inputs interrupts execution^39,41^. Furthermore, the CIS is highly predictive of the exact moment of reach initiation^27^. A similar CIS is observed in recurrent networks trained to produce reach-related muscle activity^37^. In such networks, the CIS relates mechanistically to movement triggering: it moves the neural state to a region with strong local dynamics. Working within this paradigm, the holistic and sequential hypotheses make clear predictions. There should be a monophasic CIS if compound reaches are generated holistically and a biphasic CIS if they are generated sequentially.

The CIS occurs in dimensions nearly orthogonal to those capturing reach-specific activity^27,28^. To isolate the CIS, we employed demixed principal component analysis^45,46^ to identify dimensions where activity was time-varying but largely independent of reach direction^27^. There were two such condition-independent (CI) dimensions (88% and 82% condition-invariant structure, monkey B and H). We begin by considering the projection onto one such dimension (Fig 6., *top sub-panels*). Consistent with prior findings, during single reaches there was a large change in the neural state beginning ~150 ms before movement onset and peaking around movement onset (Fig. 6a,d). There were two peaks during delayed double-reaches (Fig. 6b,e) and also during compound reaches (Fig. 6c,f).

**Fig. 6.**
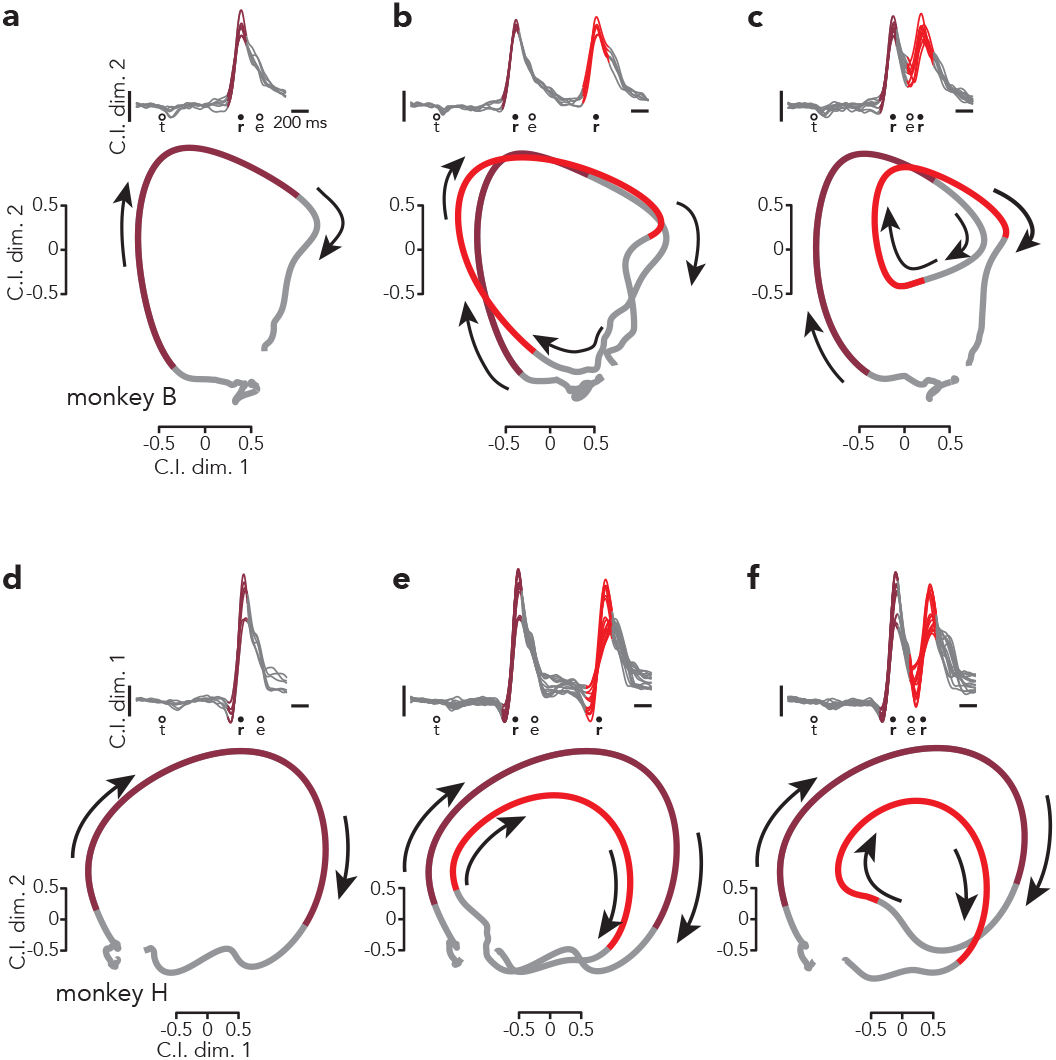
Evolution of the condition-invariant signal. **a,** *Top*: Projections of population activity from all single-reach conditions onto one condition-invariant dimension (the second, which best captured the rapid rise). The colored region of each trace highlights peri-reach activity (150 ms prior reach onset to 150 ms prior to reach end). *Bottom*: State-space projections of the same activity onto two condition-invariant dimensions. Because the projection is similar for all conditions, the average is shown to simplify presentation. As above, trace color highlights peri-reach activity. Arrows indicate the temporal evolution of activity. **b,** Same as **a** but for delayed double-reaches. The two colored regions indicate peri-reach activity for the two reaches. **c,** Same as **b** but for compound reaches. **d-f,** Same as **a-c** but for monkey H.

Prior studies have found at least two CI dimensions during reaching^27,28^. Yet visualization has typically concentrated on one dimension, either plotted versus time or as part of a three-dimensional state space where the other two dimensions capture condition-specific structure. Because we wish to concentrate on predictions regarding the CIS, we ignore that condition-specific structure and plot neural activity in a state-space spanned by two CI dimensions (bottom of panels in Fig. 6). To aid visualization, we plot a single condition-averaged trajectory, with colored portions indicating peri-reach times. Activity in the CIS dimension returns, after a reach, close to its location before the reach and thus traces out a loop (Fig. 6a,d). We stress that this loop should not be interpreted as related to the dynamics that describe condition-specific activity, which possess a rotational component^22,37^. Condition-specific dynamics occur in orthogonal dimensions^27,28^ and describe the different trajectories across many conditions. The looping trajectory in Figure 6 is similar across all conditions and is thus not evidence for a flow-field with a rotational component (indeed in models it is produced by an input rather than by network dynamics).

During single reaches (Fig. 6a,d) the looping CIS trajectory was traversed once. There were two loops during delayed double-reaches (Fig 6b,e). Because there was time for the trajectory to relax to baseline between reaches, the second loop simply began from the same region as the first. During compound reaches, the trajectory also displayed two loops (Fig 6c,f), with the second loop beginning before the first had fully relaxed. This timing lay on a continuum with that observed for delayed double-reaches with different pause durations (Extended Data Fig. 9). Thus, the CIS displays the structure predicted by the sequential hypothesis during both delayed double-reaches and during compound reaches.

### An RNN that generates compound reaches

The empirical results above universally agree with the predictions of the sequential hypothesis. We found this surprising. Not only did we expect that motor cortex would holistically generate compound reaches, it seemed implausible that it could do otherwise. Given the rapid pace of execution, the sequential strategy requires that preparation for the second reach occur during execution of the first.

That overlap is not necessarily surprising in and of itself. It is known that preparatory activity can overlap execution-related activity, both as movements are initiated (the former wanes as the latter declines^21^) and during corrections^38^. However, in all such cases the purpose of preparatory activity is proposed to be a near-immediate impact on execution-related activity. This is consistent with the proposal that a mechanistic purpose of preparatory activity is to seed execution-related activity^22,23,47^. How can preparatory activity do so for the second reach without disrupting the ongoing first reach?

To address this question, we trained a recurrent neural network to produce patterns of reach-related muscle activity (Fig. 7). As in Sussillo et al.^37^, the network received two input types. The first was a three-dimensional preparatory input that specified reach identity (*light gray traces*). The second was a condition-independent ‘go cue’ that indicated when the reach should occur (*dark gray trace*). The network then had to produce the target muscle-activity pattern for that reach (*purple traces*). We trained our network to generate two reaches on each trial, driven by two bouts of preparatory inputs and two go cues. The separation between go cues was variable and could be relatively long (Fig. 7a) or as short as 300 ms (Fig. 7b). In the latter case, the second set of preparatory inputs arrived while the network was still generating muscle activity for the first reach.

**Fig. 7.**
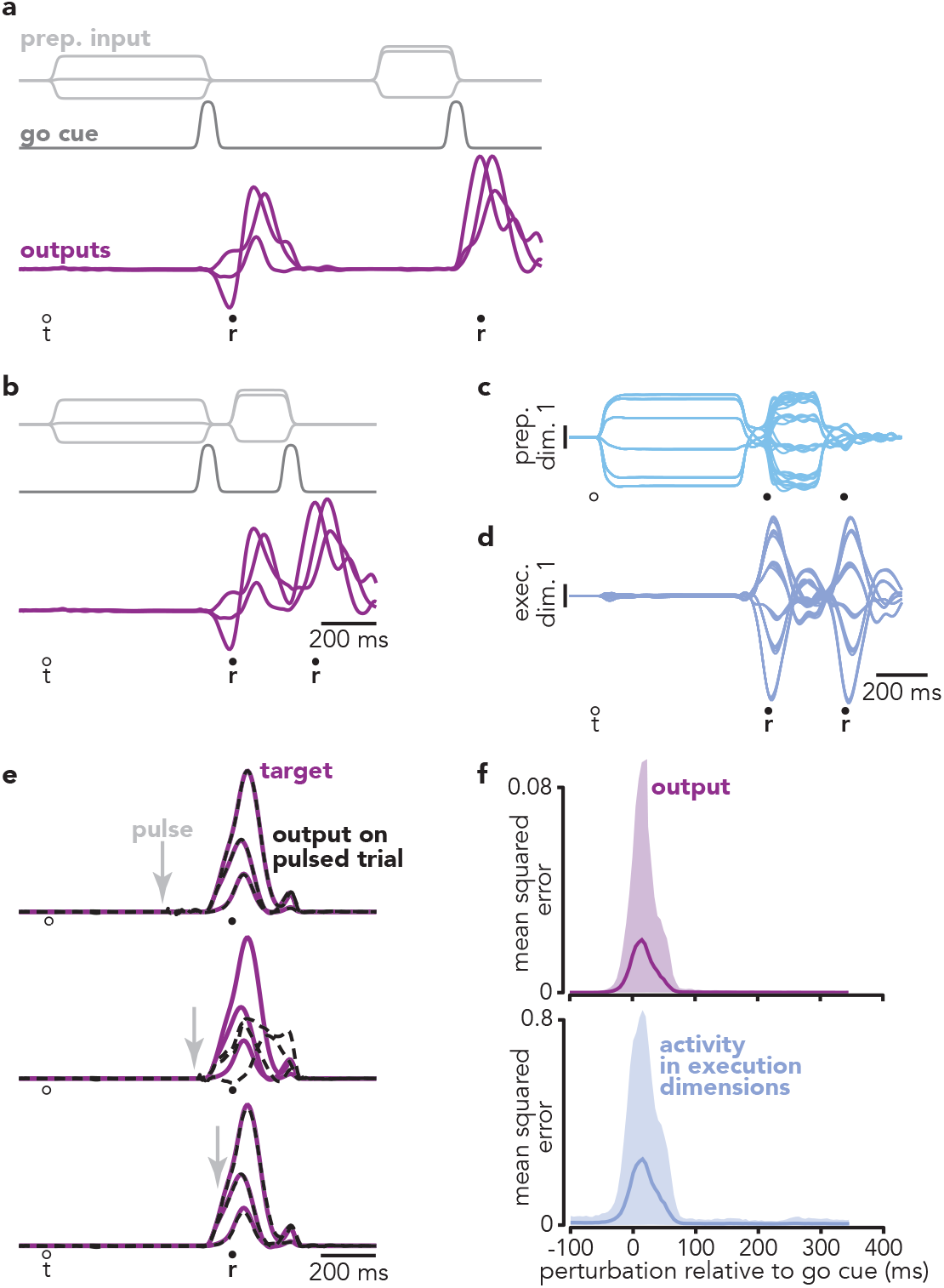
A recurrent network trained to generate reach sequences. **a,** On each trial, the network was required to produce two ‘blocks’ of output (*purple traces*), each consisting of the empirical activity of six muscles (only three of which are plotted for clarity) for one reach. Muscle activity was recorded from monkey B. The network received both a three-dimensional condition-specific ‘preparatory’ input (*light gray traces*) and one-dimensional condition-invariant a go cue (*dark gray trace*). On this example trial, the network was required to produce two blocks of output (two ‘reaches’) with a long pause between. **b,** Example trial where the inputs required the network to produce two blocks of output with no pause between, analogous to generating a compound reach. **c,** Projections of network activity onto the first preparatory dimension. Each trace plots activity for one of 36 compound reach conditions. Preparatory and execution dimensions were found using the same method applied to the neural data. **d,** Projections of network activity onto the first execution dimension. **e,** Impact of perturbing pulses, delivered to the inputs that normally convey preparatory signals. Each subpanel plots the target (*purple*) and actual (*black dashed*) output. Three (of six) output dimensions are shown for simplicity. Arrows indicate the time when the perturbation was delivered. Same time-scale as **a** and **b**. **f,** *Top*: summary of the impact of disruptive pulses on network output as a function of perturbation time. Mean squared error (between normal and actual network output) was calculated over a 200 ms window beginning at perturbation onset. Shaded region indicates the range into which 95% of effects fell. *Bottom*: same but for the impact on activity in the execution dimensions.

Our choice of inputs required the network to employ the sequential strategy – the network did not ‘know’ what the second reach would be until the arrival of the second preparatory input. We expected this scenario might be challenging and that the network might fail or adopt an idiosyncratic strategy. In fact, the network readily produced the target patterns of muscle activity across the full range of go cue separations (R^2^ > 0.99). This ability did not depend upon a fundamentally different strategy for brief go cue separations (see below). This implies that the network was able to absorb the second set of preparatory inputs, setting the stage for the second reach, without disrupting ongoing execution of the first reach. Plotting projections of network activity onto preparatory and execution dimensions (Fig. 7c,d) revealed that the second bout of preparatory activity did indeed overlap with execution-related activity for the first reach. Despite that overlap, the first-reach output was generated just as it would have been without a second reach.

To determine how the network accomplished this, we probed the link between preparation-related dimensions and network output. Because preparatory dimensions are ‘muscle null’, their influence on the output occurs via the rest of the network and is thus amenable to modulation. To explore that modulation, we employed brief input pulses that transiently perturbed activity in the preparatory dimensions. This allowed us to probe at what times, relative to the go cue, preparatory activity influences network output. Modulation was strong and temporally specific. Perturbations that coincided with the go cue disrupted network output (Fig. 7e, *middle*) while earlier or later perturbations had no effect. As a result, the influence of preparatory activity on network output was restricted to a narrow window around the time of the go cue (Fig. 7f). Critically, that influence was negligible by ~80 ms after the go cue (Fig. 7f, *top*), as was the influence on execution-related dimensions (Fig. 7f, *bottom*). Thus, not only is preparatory activity muscle-null, it is ‘dynamically null’ except during a brief window around the go cue. This allowed the network to develop preparatory activity appropriate for the second reach while muscle activity for the first reach was still being generated. The second bout of preparatory activity had an impact only at the appropriate time: after the second go cue was given.

### Comparison of neural and network strategies

Many of the basic features of the empirical data were replicated by the network model. Like empirical neurons (Fig. 8a), model units exhibited complex responses that were typically a mixture of preparation-related and execution-related activity (Fig. 8b). Despite such mixing at the single-unit level, preparatory and execution-related activity in the model occupied orthogonal subspaces, making it possible to plot projections that captured nearly pure preparatory (Fig. 7c) or execution-related (Fig. 7d) activity. Such orthogonality is noteworthy because it is a consistent feature of empirical data, but is not typically observed in networks trained to produce only single reaches^36,37^.

**Fig. 8.**
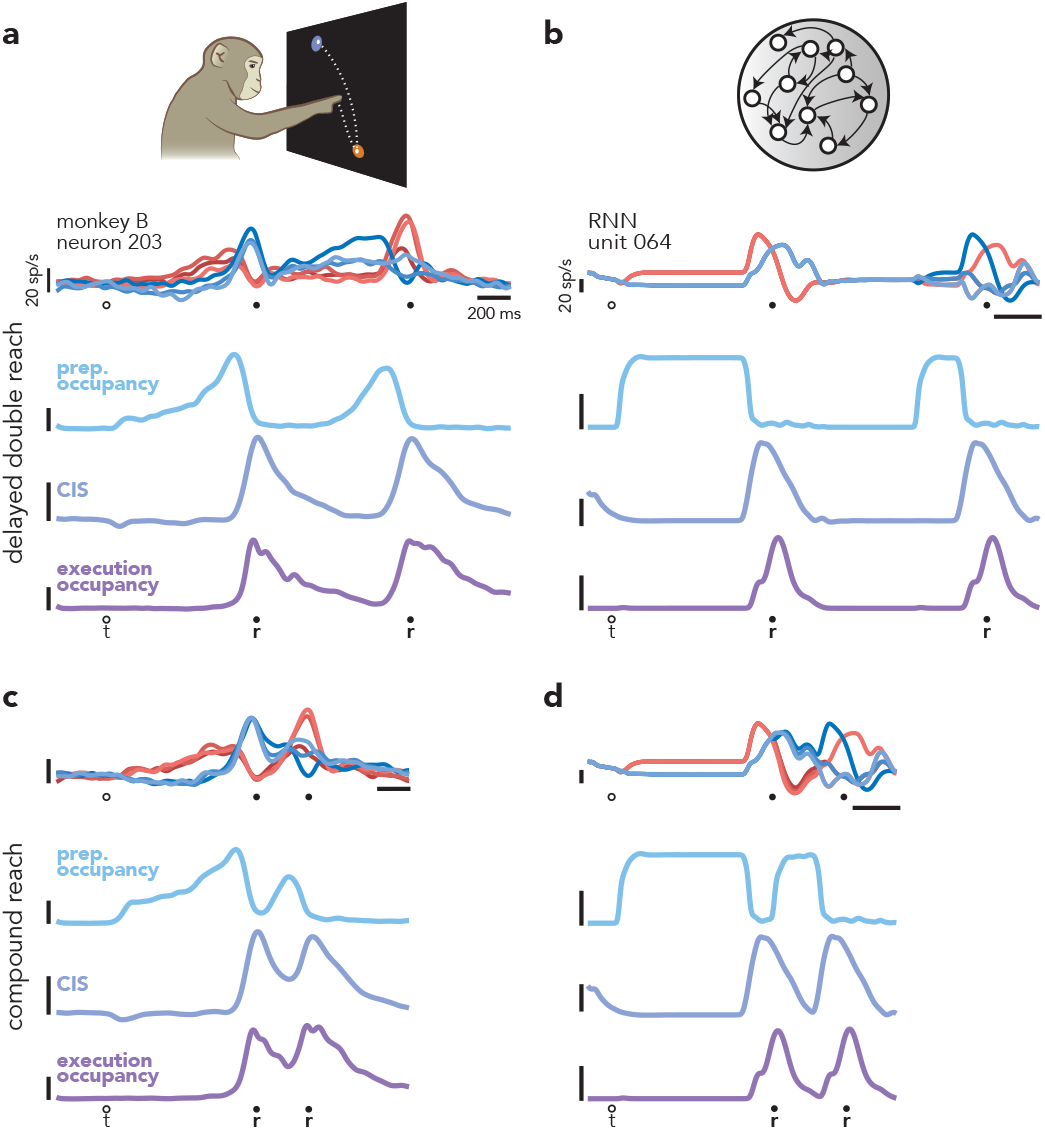
Similarities between motor cortical activity and network activity. **a,** Summary of motor cortical activity during delayed double-reach conditions. Although preparatory-, triggering-, and execution-related signals are mixed at the level of single neurons (*top*), within the population, these signals occur in orthogonal dimensions. Prior to each reach, preparatory dimensions become occupied first (*light blue trace*), then preparatory occupancy falls ~150 ms before reach onset. At approximately the same time, a condition-invariant signal (*violet trace*) occurs and occupancy of execution dimensions (*dark purple trace*) increases. This same pattern is repeated prior to the second reach. Vertical scale is arbitrary but is preserved in panel **c**. **b,** A similar sequence of events occurred in the network, which was trained to produce the empirical patterns of muscle activity from monkey B. The most notable difference between motor cortex and the network is the temporal envelope of preparatory occupancy. For the network, this results from the envelope of the network inputs; a more realistic pattern could be produced simply by altering that envelope, with essentially no change in network output. **c,** Motor cortex empirically generated compound reaches by simply repeating the sequence of events that generated delayed double-reaches. However, the second instance of the prepare-trigger-execute motif occurred shortly after the first. **d,** As was true in the empirical data, the network produced compound reaches using the same sequence of computations that produced delayed double-reaches.

The orthogonality of subspaces makes it possible to compute preparatory- and movement-subspace occupancy, just as we had for the data. Doing so reveals that, for both model and data, the sequential strategy was employed both when there was considerable temporal separation between reaches (Fig. 8a,b) and when reaches occurred in rapid succession (Fig. 8c,d). For both model and data, a given reach involved a stereotyped set of events: preparatory-subspace occupancy increased, the CIS ‘triggering’ event occurred, and execution occupancy immediately followed. Sequences of two reaches simply involved this set of events occurring twice. When the second reach occurred immediately after the first, the second bout of preparation was briefer and overlapped execution of the first reach.

## Discussion

### The Sequential Strategy Can Produce Rapid Movement Sequences

Motor sequences have been studied for over one hundred years^48–50^, yet it has remained unclear how low-level computations contribute to generating sequential behavior. Does producing a rapidly unfolding motor sequence require individual elements to be fused into larger, cohesive units? Or can motor cortex leverage a unified strategy to generate sequences at different paces?

In our task, compound reaches potentially appeared to be holistically generated at the level of the muscles: muscle activity formed one continuous pattern spanning the full compound reach. Nevertheless, motor cortical activity revealed that component reaches were prepared and executed sequentially. Compound reaches involved two epochs of preparatory activity, each of which reflected the identity of the immediately pending reach rather than the full sequence. Compound reaches also involved two peaks of the condition-invariant signal, which is hypothesized to reflect movement triggering. Execution-subspace activity reflected the reach being currently generated rather than the full sequence – e.g., it was very similar during the first reach regardless of the presence or identity of the next reach.

The ability to separate preparation-related and execution-related activity was central to many of the above findings and also revealed how the rapid pace of compound reaches was achieved. That pace depended not on the preparation of multiple reaches prior to initiation, but upon simultaneously engaging preparation and execution. That temporal overlap allowed the second reach to be prepared while the first was still unfolding. Related forms of overlap have been recently observed: preparatory and execution-subspace activity overlap around movement onset^21^ and during externally driven corrections^38^. However, in all such cases the presumed goal of preparatory-subspace activity was to immediately impact motor output. We were initially surprised that preparatory subspace activity could develop without impacting either execution-subspace activity or motor output. However, a neural network trained to generate sequential reaches readily utilized this strategy. In doing so, it adopted internal activity that displayed the empirically observed features described above (e.g., two bouts of preparation, triggering, and execution).

We stress that our results do not prove that it is impossible to holistically generate a multi-element movement. Our goal was not to determine whether the holistic strategy is possible, but whether a holistic strategy is essential to the rapid production of motor sequences. Based on our findings, it is clear that a holistic strategy is not necessary for motor cortex to generate rapid sequences, nor does it emerge from familiarity with a particular sequence^3,15,29^. Our monkeys had extensive practice with compound movements and performed them so rapidly that muscle activity merged into a continuous pattern, yet the sequential strategy was employed at the level of motor cortex. Nevertheless, it is quite possible that a holistic strategy could be encouraged by specific training approaches, perhaps by training on compound reaches alone (but never their component reaches) or by restricting sensory cues^51^. As will be discussed below, a trivial version of the holistic strategy -- replacing two movements with one that achieves a similar goal – is almost certainly possible.

### Extending the Prepare-Trigger-Execute Model

The prepare-trigger-execute paradigm was developed in the context of single reaches and how they could be generated by networks with strong internal dynamics. Whether this paradigm is applicable to motor sequences is test of the framework’s utility: do the data agree with self-consistent sets of predictions made by competing hypotheses developed within that paradigm? This was indeed the case, and it is worth emphasizing that this did not have to be true. The data could have agreed with some predictions of one hypothesis and some of the other, or with neither. Mixed or difficult-to-interpret results would have indicated that the paradigm needed to be reevaluated. Instead, in every case the data obeyed the predictions of the sequential strategy and not the holistic strategy. These results serve not only to support one hypothesis and reject the other, but also to demonstrate the utility of the paradigm used to couch the hypotheses and generate the predictions.

Our results also help extend the prepare-trigger-execute paradigm. Preparation has historically been considered a time-consuming process^47,52–55^ and thus to contribute to the longer time between sequence elements at chunk boundaries^56,57^. This made it reasonable to assume that minimizing temporal separation required a holistic strategy. However, it has now been demonstrated that preparation can be remarkably swift for well-practiced movements^21,58^. The present results confirm that swift preparation is possible, allowing rapid sequences to be produced via the sequential strategy.

Our results also agree with recent findings showing or suggesting that preparatory activity can occur during movement. Ames et al.^38^ found that preparatory dimensions become occupied during movement when a jumping target required an immediate correction, and Stavisky et al.^59^ found that the response to an unexpected visual perturbation occurs first in an output-null space before flowing into an output-potent space. The present study is the first to show that the preparatory subspace can be occupied in anticipation of a pending movement without causing a change in the present movement. This presumably depends upon the orthogonality of preparation and execution-related dimensions. Such orthogonality naturally appeared when we pushed networks at a pace that required preparation and execution to overlap. In contrast, the networks in^37^ were never trained to perform reach sequences and did not show orthogonal preparation and execution. Thus, the need to develop preparatory activity despite ongoing movement provides a possible explanation for a consistent feature of the empirical population response. Additionally, the orthogonality of preparatory and execution dimensions may relate to optimal control of preparatory activity^60^.

### Generating Sequences of Actions

A consequence of the sequential strategy is that, in caudal PMd and M1, activity before sequence onset reflects only the first reach. Yet upstream areas must presumably look further ahead. Previous work demonstrates that delay-period activity in the supplementary motor area (SMA) reflects the identity of upcoming two-element^61^ and three-element^62^ sequences. A similar segregation has also been found in human subjects performing a key-press task; while there is no evidence for sequence-specific selectivity in the M1 BOLD signal, such selectivity is present in upstream areas^16–18^. Using a cycling task, which shares some features with sequence tasks, Russo et al.^63^ found that M1 activity reflects only the present motor output while SMA activity distinguishes situations that will have different future outputs. When we began this study, a reasonable hypothesis was that SMA coordinates sequences – e.g., specifies which elements will be produced in what order^64^ – allowing motor cortex to generate closely spaced elements holistically. Given present results, upstream computations must instead enable the sequential strategy, which requires carefully timed preparatory and triggering inputs. These inputs likely originate not only from SMA but from many areas, both cortical^62,65–67^ and subcortical^68–70^.

Although skillfully and rapidly performing a sequence of motor elements does not depend on a holistic strategy at the level of motor cortex, there may still be cases where skilled action depends on replacing elements – i.e., learning new movements^4^. For example, when learning to sign their name, a person transitions from writing each letter deliberately to producing the entire signature in fewer, smoother strokes. Of course, the shape of individual letters changes throughout this process, as do the patterns of muscle activity that produce them. There is thus an important distinction between learning to more rapidly link elements (but preserving the elements themselves) and replacing one set of actions with an entirely new one that serves a similar high-level goal but is otherwise a different movement^64,71^. The former can be driven by a sequential strategy (with greater skill perhaps resulting from more rapid and consistent pacing of the different stages) while latter requires the holistic strategy by definition.

## Acknowledgments

We thank Y. Pavlova for excellent animal care. This work was supported by the Grossman Center for the Statistics of Mind, the Simons Foundation (M.M.C.), the McKnight Foundation (M.M.C.), NIH Director's New Innovator Award DP2 NS083037 (M.M.C.), NIH CRCNS R01NS100066 (M.M.C.), NIH 1U19NS104649 (M.M.C.), P30 EY019007 (M.M.C), the National Science Foundation (A.J.Z.), and the Kavli Foundation (M.M.C.),.

## Methods

### Subjects and Task

Subjects were two adult, male macaque monkeys (monkeys B and H). Animal protocols were approved by the Columbia University Institutional Animal Care and Use Committee (AC-AAAM2550). Experiments were controlled and data collected under computer control (Speedgoat Real-time Target Machine). During experiments, monkeys sat in a customized chair with the head restrained via a surgical implant. Stimuli were displayed on an LCD monitor in front of the monkey, and a tube dispensed juice rewards. The left arm was comfortably restrained, and the task was performed with the right arm.

Hand position was monitored using an infrared optical system (Polaris; Northern Digital) to track (~0.3 mm precision) a reflective bead temporarily affixed to the third and fourth digits. Each trial began when the monkey touched and held a central touch-point. After holding for 400-600 ms (randomized) targets appeared – one for single reach conditions and two for delayed double-reach and compound reach conditions. There were six possible target locations, arranged radially 140 mm from the touch-point. Targets were round and had to be hit with within 25 mm of their center. A condition was defined by target location(s) and whether success required a single reach, a delayed double-reach, or a compound reach. Conditions were interleaved using a block-randomized design.

For single reach conditions, a 20 mm diameter, green target appeared at one of the six locations. After a variable delay period (0-1000 ms), the target doubled in size and the central touch point disappeared. These events served as salient go cue, instructing the monkey to reach. Reaches were successful if they were initiated 100-500 ms after the go cue, lasted <500 ms, and stayed on the target for 600 ms with minimal hand motion. These requirements were conserved across all conditions.

For compound reach conditions, two targets appeared simultaneously. Target color indicated the first (green) and second (blue) targets. After a variable delay period (0-1000 ms) both targets grew in size and the touch point disappeared, providing the go cue. The monkey then captured the first target and was immediately given a juice reward. After capturing the second target (and holding for 600 ms) the monkey was given a second juice reward. During compound reach conditions, we encouraged the monkey capture the second target as quickly as possible after capturing the first. We used a speed threshold (70 mm/sec, roughly 5% of peak velocity) to determine when the first target was captured. As soon as hand velocity fell below this threshold, the second target began to shrink rapidly, such that it disappeared (and the trial was aborted) after 350 ms. Shrinking stopped when the second reach began, providing incentive to do so quickly. In practice, monkeys reached considerably faster than required, with a median pause between reaches of 119 ms (monkey G) and 137 ms (monkey H).

Delayed double-reach conditions started as for compound reaches, with a key exception. The first target was colored only in its center – the rest was white. The diameter of the colored center indicated the duration of the imposed pause between reaches. Diameters of 8 mm, 14 mm, and 18 mm indicated pauses of 600 ms, 300 ms, and 100 ms. The imposed pause began when the hand reached the first target. At that moment, the colored portion of the target began to grow at a constant rate. The monkey’s hand was required to remain within the target with minimal movement until the colored center reached the perimeter of the first target (i.e., the target was all green). Immediately after becoming all green, the first target disappeared and the second target instantly grew in size, indicating that the monkey was now permitted to reach towards the second target. A successful second reach required initiation within 500 ms (<100 ms reaction times were allowed here, as the time of the second go-cue was predictable) and then holding the final target for 600 ms. Reward was delivered after both the end of the pause between reaches and at the end of the final hold period.

### Neural and Muscle Recordings

After initial training, we performed sterile surgery to implant a head restraint and cylindrical recording chamber, which was positioned to give access to the arm area of dorsal premotor cortex (PMd) and primary motor cortex (M1). Recordings for both monkeys were performed in the left hemisphere. The task was performed with the right arm. Chamber positioning was guided by structural magnetic resonance imaging prior to implantation. We used intra-cortical microstimulation to confirm that our recordings were from the forelimb region of motor cortex (biphasic pulses, cathodal leading, 250 ms pulse width delivered at 333 Hz for a total duration of 50 ms). Microstimulation typically evoked contractions of the shoulder and upper-arm muscles, with current thresholds between 20μA-200μA, depending on location and stimulation depth (a wide range of current thresholds is expected, as we recorded in both M1 and PMd and from a variety of depths). We recorded single-neuron responses using one or more 32-channel linear-array electrodes (S-probes; Plexon) lowered into cortex using a motorized microdrive. Spikes were either sorted offline by hand (Plexon Offline Sorter) or automatically using KiloSort^72^. Spike clusters found by KiloSort were manually curated (Phy Template-gui software; Kwik Team). We recorded all well-isolated task-responsive neurons; no attempt was made to screen any response property. Spikes were smoothed with a Gaussian kernel with standard deviation of 25 ms and averaged across trials to produce peri-stimulus time histograms. Across all recorded neurons, the average minimum trial-count per condition was ~20 trials.

In dedicated sessions, we recorded electromyogram (EMG) activity using intramuscular electrodes from the following muscles: trapezius, anterior, lateral, and posterior heads of the deltoid, lateral head of the biceps, pectoralis, and brachialis. EMG signals were bandpass filtered (50-5kHz), digitized at 30 kHz, rectified, smoothed with a Gaussian kernel with standard deviation 25 ms, and averaged across trials to produce a continuous estimate of muscle activity intensity.

### Data Pre-Processing

Photodetectors (Thorlabs) were used to synchronize commands from the behavioral control software with the 60 Hz refresh rate of the display, such that the timing of visual events was known with 1 ms accuracy. Hand position was sampled at 60 Hz. For analysis on a one millisecond timescale, values were interpolated and filtered. For analysis, movement onset and offset were estimated offline based on hand speed. Movement onset (for each reach) was defined as the time that hand speed first surpassed 5% of the peak speed for that reach. Similarly, movement end was defined as the time when hand speed first dropped below 5% of the peak reach speed.

In order to provide a clear view of preparatory activity during the instructed delay period, all analyses only included trials with instructed delays >500 ms. To provide a continuous view of neural activity during the delay period, data were aligned to target onset and the onset of the first reach and concatenated. Concatenation occurred 350 ms prior to the onset of the first reach.

To compare the temporal evolution of neural activity across time and conditions, we aligned single trials to key behavioral events. For each set of conditions that shared a temporal task structure, we scaled the time-base for each trial such that the duration of the first reach and the dwell-time between reaches matched the median first reach duration and dwell-time for that set of conditions. This was done separately for single reaches, compound reaches, and delayed double-reaches. Monkey B performed delayed double-reaches with three different pause durations; the above procedure was performed separately for each. All variables of interest (firing rate, hand speed, EMG) were computed for each trial prior to alignment. Thus, alignment never alters the magnitude of these variables, it simply modifies slightly when they occur. Using an alternative alignment procedure that did not involve rescaling – aligning single-trial data to target onset, first reach onset, and second reach onset and concatenating – produced very similar results.

### Determining Sequence Selectivity

We defined sequence selectivity in the traditional way: responses that were statistically different when a reach was part of a sequence versus performed alone. First reaches provided the most natural comparison, as they were physically similar when performed alone versus as part of a sequence. We thus asked whether activity before and during first reaches differed between the single-reach and compound-reach conditions. We computed the average firing rate on each trial during two epochs: the delay period and an ‘execution-epoch’ spanning the first reach. A t-test (unpaired) was used to ask whether, for each epoch, these differed for single versus compound reaches. For the execution epoch, we employed a second comparison that avoids the concern that differences might be missed when averaging across time-points. We compared, at each millisecond, firing-rates for single versus compound reaches, and took the average absolute difference. This was summed across conditions (so that we made one comparison per neuron). We used a resampling test to determine if the summed difference was greater than expected given trial-to-trial variability. Resampling was performed by pooling all single and compound reach trials, and drawing two new ‘conditions’ (sampling with replacement). Differences were considered significant at the 0.001 level if the true difference was greater than 1000 resampled differences.

### Identifying Preparatory and Execution Dimensions

As in previous work^21,22^ we employed two additional preprocessing steps prior to dimensionality reduction. First, we soft-normalized each neuron’s firing rate. The normalization factor was equal to that neuron’s firing rate range plus five spikes/s. This encouraged dimensionality reduction to capture the responses of all neurons, rather than just high firing-rate neurons. Second, within each set of conditions of the same duration (e.g., single reaches, compound reaches) we mean-centered the firing rate of each neuron, so that its average rate (across conditions) was zero at all times. This step ensures that dimensionality reduction focuses on dimensions where activity is selective across conditions. Dimensions capturing condition-invariant activity^27,28,39,40^ were identified separately (see below). We employed a recently developed dimensionality reduction approach^21,36,38^, which leverages the fact that motor cortex population activity occupies nearly orthogonal subspaces during preparation and execution^36^. Importantly, this remains true even for preparatory events that occur outside an instructed delay period^21,38^. Identifying those dimensions requires two sets of data: one where activity reflects only preparation and another where activity reflects only execution.

Preparation-only activity was obtained using task epochs where the hand was stationary but preparation was presumed to occur. Most basically this included a ‘delay-period’ epoch from 700 – 150 ms prior to the onset of single reaches. We similarly employed the delay-period epoch before compound reaches and delayed double-reaches. For the delayed double-reaches, we used trials with a 600 ms inter-reach pause (which was experienced by both monkeys). These same trials provided an additional ‘second-reach preparatory’ epoch: 300 ms – 150 ms prior to the onset of the second reach. We included this epoch because second reaches had trajectories that were different from first reaches (they moved from one peripheral target to the other, rather than from the center out) and we wished preparatory dimensions to capture preparation for both first and second reaches. To be conservative, we did not use a second-reach epoch before compound reaches; one of the competing hypotheses predicted no preparatory activity at that time, and we wished to avoid any concern that our method might ‘build it in.’

Execution-only activity was obtained from a peri-reach epoch, −50 – 500 ms relative to reach onset. We used data only for reaches where it was unlikely that preparation overlapped execution. This included single reaches, both reaches for delayed double-reaches with a 600 ms pause (the longest pause we used), and the second reach of compound reaches. For all these reaches, any subsequent reach (either second reaches or the return to the center) is some distance in the future, such that any preparation for the next movement likely occurs after completion of the present reach.

Using activity from the above times, we constructed two matrices: *P* ∈ *R^N x T^_prep_* which holds the neural activity from preparatory epochs and *E* ∈ *R^N x T^_exec_* which holds activity from the execution epochs. *N* is the number of recorded neurons, *T_prep_* is the total number of times across all conditions and all preparatory epochs as defined above, and *T_exec_* is the total number of times across all conditions and all execution epochs as defined above. We sought a set of preparatory dimensions *W_prep_* that maximally capture the variance in *P*, and an orthogonal set of dimensions *W_exec_* that maximally capture the variance of *E*. To do so, we computed covariance matrices *C_prep_* = *cov(P)* and *C_exec._*=*cov(E)* and used numerical optimization to find:

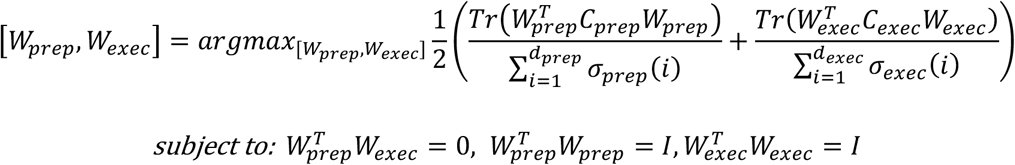

where *σ_prep_(i)* is the *i^th^* singular value of *C_prep_* and *σ_exec(i)_* is the *i^th^* singular value of *C_exec_*. *Tr(·)* is the matrix trace operator. 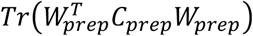 is the variance captured, across all preparatory epochs, by the preparatory dimensions. 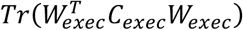 is the variance captured, across all execution epochs, by the execution dimensions. The optimization objective is normalized (by the singular values) to make it insensitive to differences in total preparatory versus execution variance. Using a dimensionality of 20, *W_prep_* and *W_exec_* captured 78% and 68% of the variance in all preparatory epochs for monkey B and monkey H, respectively and 68% and 70% of the variance in all execution epochs for monkey B and monkey H, respectively.

Some analyses, in particular those involving visualization, involved focusing on a subset of the dimensions within *W_prep_*. To do so, we found *W_prep_rotate_*, a set of dimensions spanning the same subspace but ordered so that the first dimension captures the most variance and so forth (as for PCA). To avoid this rotation being biased towards one type of reach, it was based on preparatory epoch activity from an equal number of times before first versus second reaches. This encouraged the first two dimensions to capture preparation-related activity before both first and second reaches (it did not ensure equal variance for both, but encouraged parity that if indeed preparatory activity was of similar magnitude for both). To find *W_prep_rotate_*, we projected a 200 ms window of activity before all six single reaches (starting 300 ms before reach onset) and a 200 ms window of activity before six delayed double-reaches (starting 300 ms before the onset of second reach) onto *W_prep_*, yielding a matrix *Q* ∈ *R*^20 × 12*T*^, where *T* was equal to 200. We performed PCA on *Q*, yielding a rotation matrix *B* ∈ *R*^20 × 20^. *W_prep_rotate_*was then *W_prep_B*. All population analyses were performed by projecting population data onto *W_prep_rotate_*. This rotation of the basis set was important when focusing on a subset of dimensions (one wishes to prioritize variance captured) but had no impact on analyses that employed all dimensions (e.g., *W_prep_* and *W_prep_rotate_* capture the same total variance).

### Subspace occupancy

We calculated occupancy as described in^21,36^. Below we describe how we computed preparatory occupancy. Execution occupancy was computed analogously. For a given time *t* and condition *Φ*, the projection of the population response onto the preparatory dimensions is: *x^prep^*(*t*, *ϕ*) = *W_prep_rotate_* ***r***(*t*, *ϕ*), where ***r***(*t*, *ϕ*) is a vector containing response of each neuron for time *t* and condition *ϕ*. To measure subspace occupancy, we calculated:

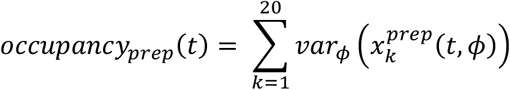

Where *var_ϕ_* indicates taking the variance across conditions and 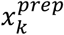 is the *k^th^* element of *x^prep^*. We employed a resampling procedure to estimate sampling error. We created 1000 surrogate neural populations by redrawing, with replacement, neurons from the original population. For each, we computed the preparatory dimensions and subspace occupancy, yielding a distribution of occupancies. That distribution was used to compute error bars and for assessing statistical significance. For example, it allowed us to ask whether preparatory subspace occupancy during the first compound reach was significantly higher than that during single reaches.

To compare preparatory occupancies between single reaches and compound reaches, we asked within a particular window (200 ms window ending with the onset of the second reach), how frequently (across surrogate populations) was preparatory occupancy higher for compound reaches than for single reaches. If compound reach occupancy was higher in 95% of comparisons, this would yield a p-value of 0.05.

### Identifying Trigger Dimensions

We used dPCA^45,46^ to identify neural dimensions where activity varies primarily with time and not condition^27^. dPCA was applied to a matrix of firing rates, *A* ∈ *R^CT x N^*, where *N* is the number of neurons, *C* is the number of conditions, and *T* is the number of time-points per condition. The data in *A* was taken from a 300 ms window centered on reach onset. Data was included for all of the reaches from single-, compound-, and delayed double-reach-conditions – i.e., all reach directions and reach types. dPCA leverages lables assigned to each row of *A* that identify the condition and time. dPCA seeks a matrix *W_dPCA_* ∈ *R^N x k^*, where *k* is specified (we chose *k* to be 8), that produces a projection *X* = *AW_dPCA_*. Each column of *W_dPCA_* is a dimension and each column of *X* is thus a projection of the population response onto that dimension. Like PCA, dPCA optimizes dimensions to capture variance, such that 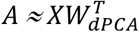. Unlike PCA, dPCA further optimizes *W_dPCA_* such that each column of *X* covaries strongly with only one label (i.e., time or condition). Our analysis of the condition-invariant signal employed those columns where activity varied primarily with time. We observed very similar results if compound reach conditions were omitted when finding *W_dPCA_*.

### Training a Recurrent Neural Network to Generate Sequences of EMG

To determine whether a recurrent neural network (RNN) could readily use the sequential strategy, we trained an RNN to reproduce the empirical patterns of muscle activity, largely following the procedure outlined in^37^. We used a network with dynamics:

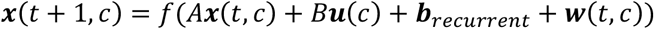

where ***x*** is the network state for time *t* and condition *c*. The function *f* was a hyperbolic tangent linking each unit’s inputs to its firing rate, *A**x*** captures the influence of network activity on itself via connection weights *A*, *B**u*** represents the external inputs, *b_recurrent_* is a vector of offset biases, and the random vector ***w*** ~ *N*(0,*σ_w_I*) adds a small amount of noise. We set *σ_w_* to be 5×10^−4^. Network output was a linear readout of ***x***:

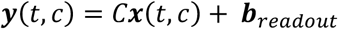

The parameters *A, B, C,* ***b***_*recurrent*_, and ***b***_*readout*_ were optimized to minimize a loss function composed of the difference between the network output, ***y*** and a target, ***y***_*targ*_ and a number of regularization terms (see below). The target output was based upon EMG recorded from six muscles from monkey B during all six individual reach conditions. We created 36 ‘conditions’ – each a two-reach combination – that the network had to perform. Each condition was generated by concatenating muscle activity from two reaches, separated by an interval where the target output was 0. On each ‘trial’ the network performed one condition, with a random value of that interval (0 – 600 ms). Thus, the model was asked to produce six-dimensional muscle activity for all 36 possible two-reach combinations, and to do so across a wide range of delays.

The loss function optimized during training was:

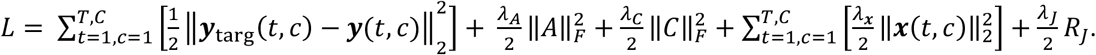

The first term is the error between the network output and the target. All other terms serve to regularize the network solution. The second and third terms penalize large recurrent and output weights, respectively. The fourth term penalizes large firing rates, and the final term discourages complex dynamics^37^:

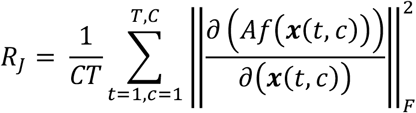

This final term was not necessary for the network to successfully learn to produce the target output, nor the general solution it found (i.e. placing preparatory, triggering, and execution activity in largely orthogonal sets of dimensions). However, as in^37^, regularization in general and the inclusion of *R_J_* in particular encouraged network solutions that resembled, at the population and single-neuron level, those observed empirically. Regularization coefficients were chosen to be: *λ_A_* = 10^−7^, *λ_C_* = 10^−7^, *λ_**x**_* = 10^−8^, and *λ_J_* = 5 × 10^−5^. The RNN was composed of 100 units, received 3 condition-specific inputs, and a single condition-independent ‘go-cue’. In order to ensure that the network was truly producing an output in response to the go-cue (and not implicitly time-locking to the start of simulation) we used a variable delay between condition-specific input onset and go-cue onset; this delay varied randomly between 200 and 600 ms. The matrices *A, B,* and *C* were initialized as random orthonormal matrices, and the network was trained using TensorFlow’s Adam optimizer.

We asked how the network responded to disruptive pulses delivered around the time of the go-cue. Each pulse was delivered via the network inputs for 30 ms and was a scaled version of the input vector associated with one of the non-tested (i.e. off-target) reach directions. The magnitude of each pulse vector was between 0.5 and 3 times that of the original input vector. Each time-point in Fig.7f was probed with 1000 pulses.

---

**Extended Data Fig. 1.**
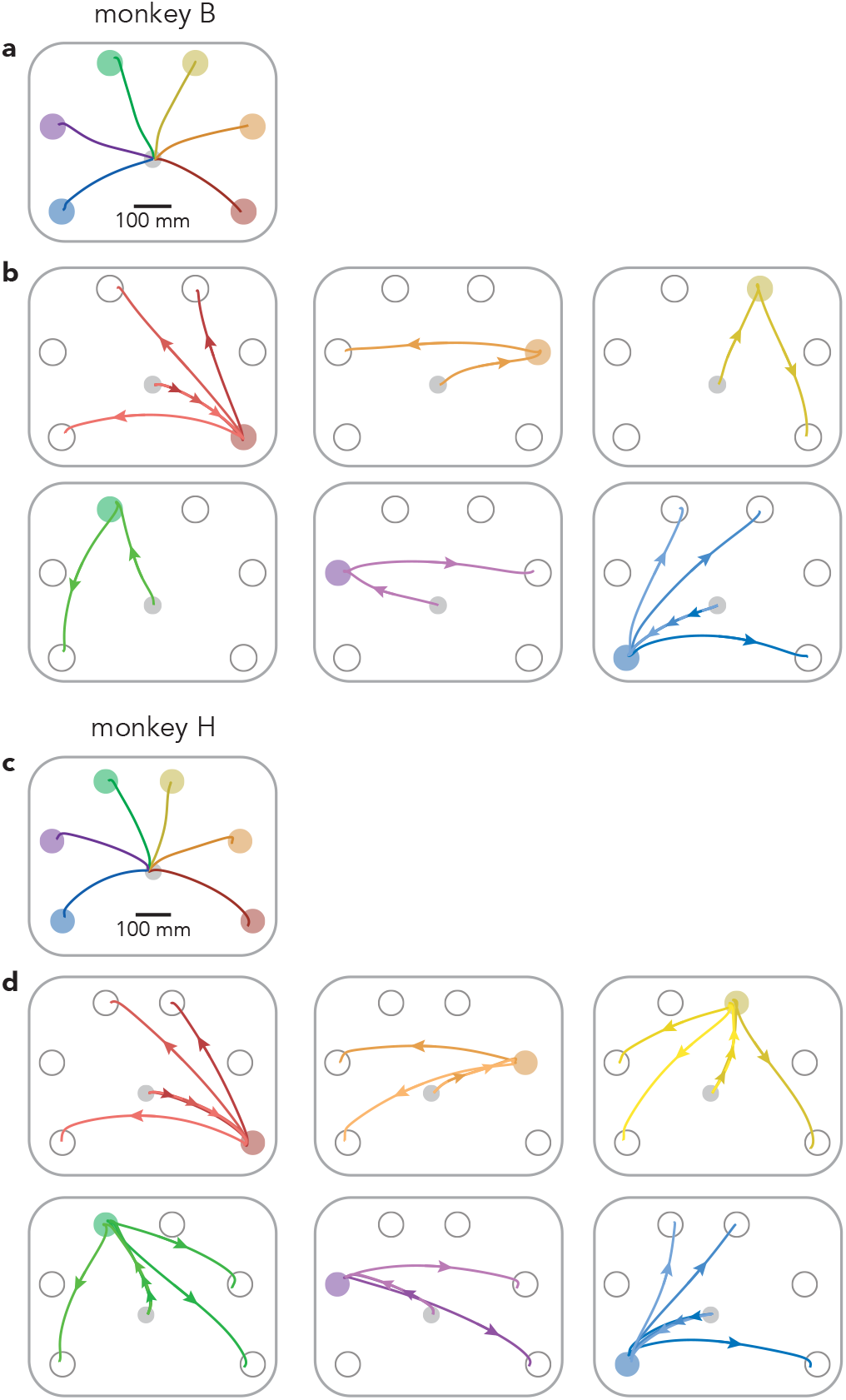
Single and compound-reach conditions. **a,** Reach paths for single reach conditions (same as in Fig. 1a) for Monkey B. Paths are averaged across all trials and sessions. **b,** Reach paths for all compound reach conditions performed by monkey B (a superset of those in Fig. 1a). Most first-target locations were used for only one compound reach condition, to maintain a reasonable trial-counts. This was necessary because monkey B performed delayed double-reach conditions using three instructed pauses, which added to the total number of conditions performed. For similar reasons, delayed double-reach conditions employed a subset of the two-target combinations (those in *red* and *blue*) that were employed during compound reaches. **c,** Same as **a,** but for monkey H. For monkey H, the bottom-right and top-right targets were shifted slightly to the right and left (respectively) compared to the locations used for monkey B. This shift was necessary to prevent the animal’s arm from blocking sight of the second target during certain conditions (the two monkeys were of different sizes and employed slightly different postures when reaching). **d,** Same as **b,** but for monkey H. Monkey H performed a greater number of compound reach conditions than monkey B.

**Extended Data Fig. 2.**
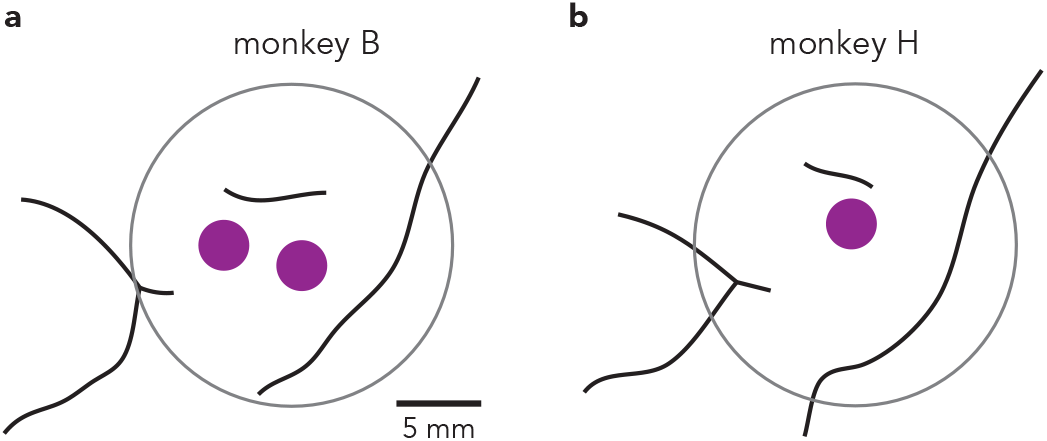
Recording locations. **a,** Location of recording chamber (*grey*) and recording area (*purple*) relative to arcuate sulcus, pre-central dimple, and central sulcus (*black*, left to right). **b,** Same as **a** but for Monkey H.

**Extended Data Fig. 3.**
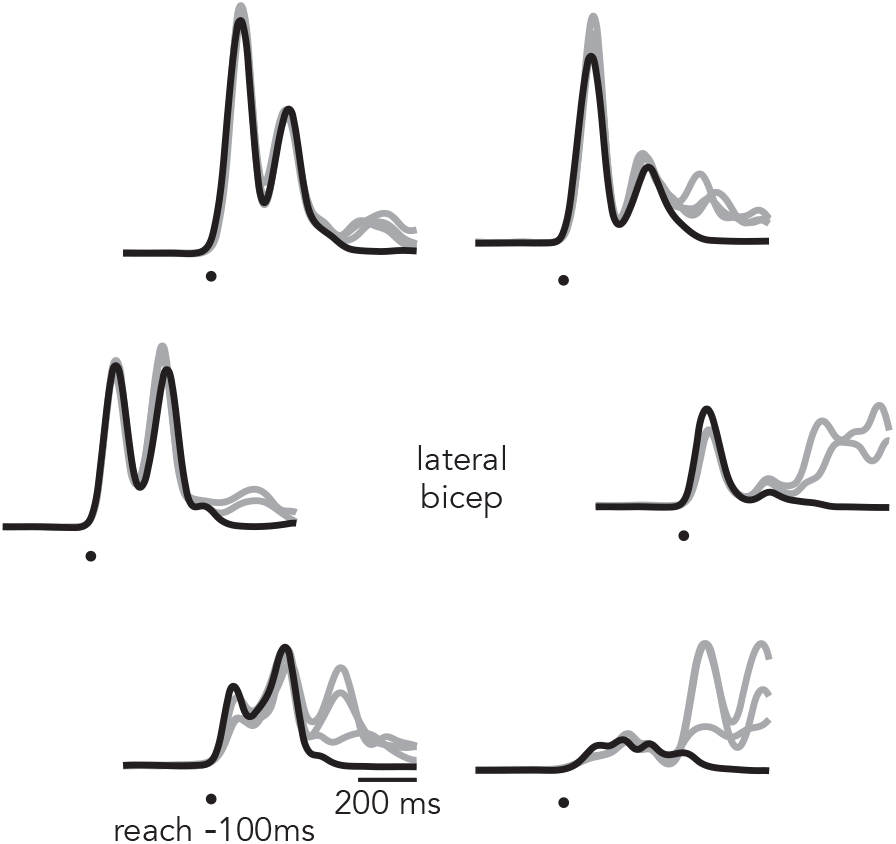
Additional example EMG. Mean EMG for single (*black*) and compound (*grey*) reach conditions, Monkey H.

**Extended Data Fig. 4.**
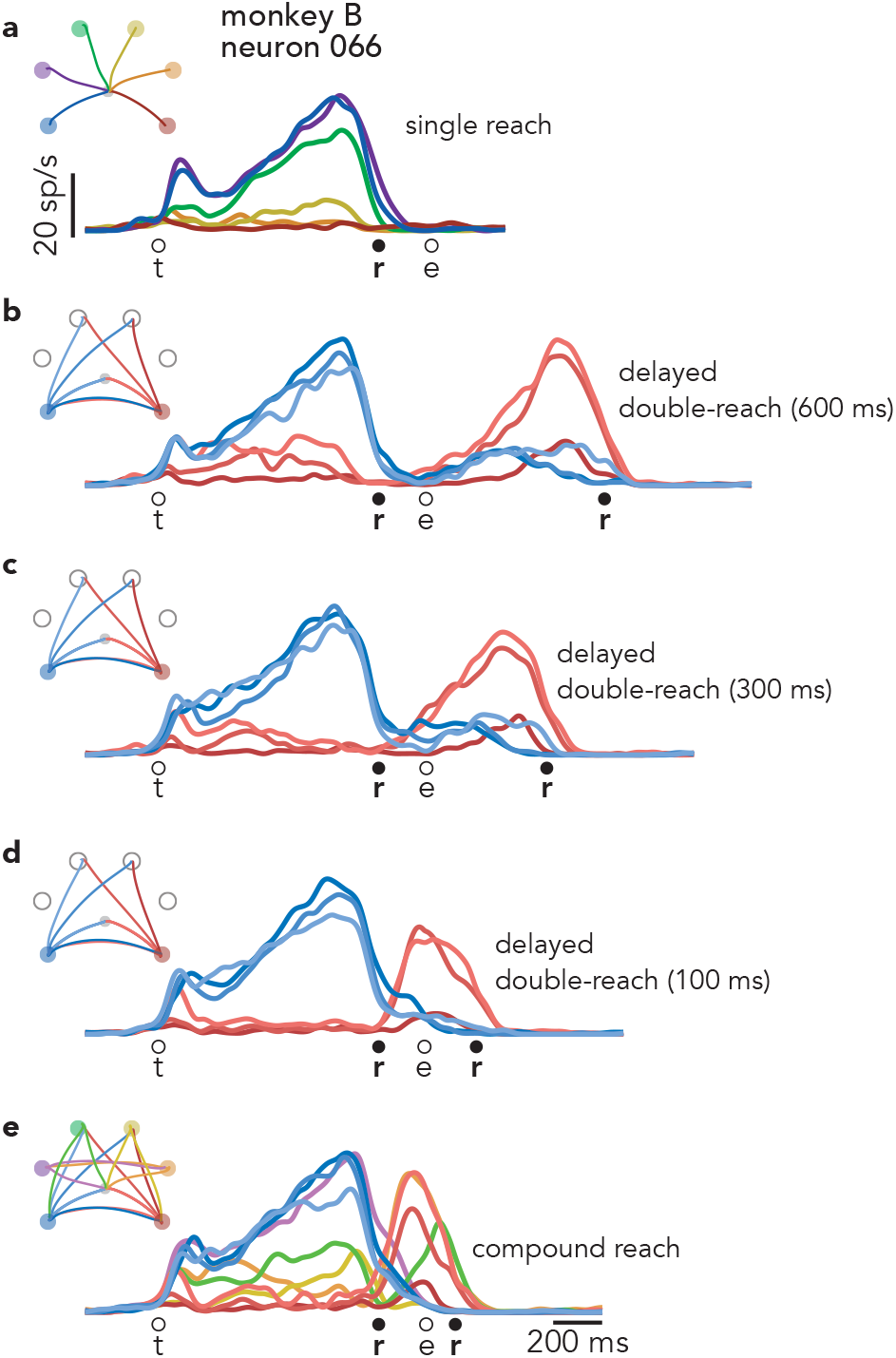
Activity of neuron 066 (Monkey B) across all conditions. **a,** Response during single-reach conditions. This panel plots the same neural data as in Fig. 3a, but for all single reaches. **b,** Response during delayed double-reaches with a 600 ms instructed pause. This panel plots the same neural data as in Fig. 3b. **c,** Response during delayed double-reaches with a 300 ms instructed pause. **d,** Response during delayed double-reaches with a 100 ms instructed pause. **e,** Response during all compound reach conditions. The traces in this panel are a superset of those in Fig. 3c.

**Extended Data Fig. 5.**
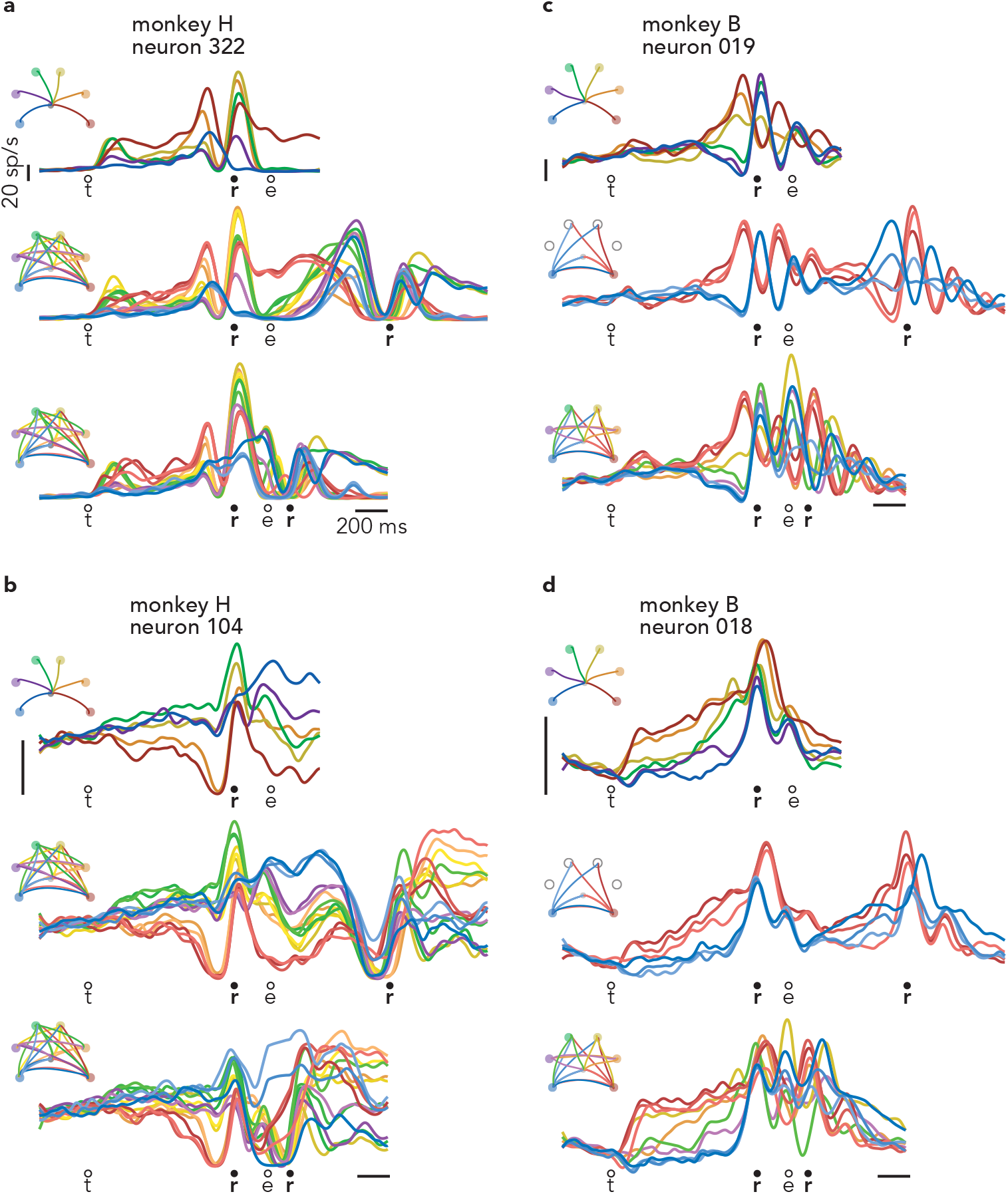
Responses of four exemplar motor cortex neurons. **a,** Response of neuron 322, recorded from Monkey H. These are the same neural data as in Fig. 3d-f, but are plotted here for all single reaches and all two-reach combinations. **b-d,** Responses of three additional example neurons. As is typical, these neurons are active during both the delay and execution-epochs of single reaches. It is thus difficult to determine via inspection whether there is a second bout of preparation during compound reaches.

**Extended Data Fig. 6.**
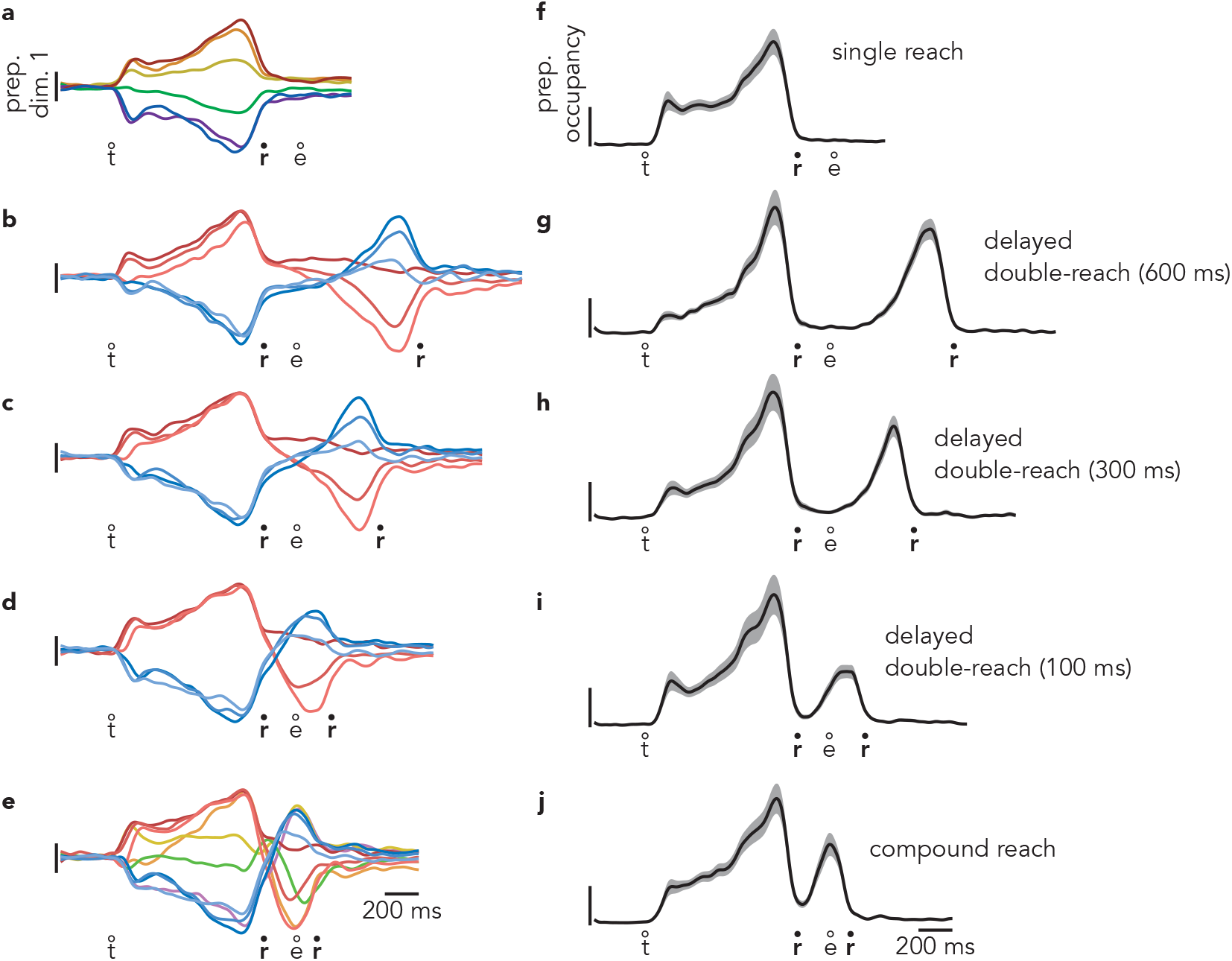
Time-course of activity in preparatory dimensions during all conditions, monkey B. **a,** Projections of population activity during single reach conditions onto the first preparatory dimension. This panel plots the same data shown in Fig. 4a. **b,** Activity in the first preparatory dimension during delayed double-reach conditions with a 600 ms instructed pause. This panel plots the same data shown in Fig. 4b. **c,** Activity in the first preparatory dimension during delayed double-reach conditions with a 300 ms instructed pause. **d,** Activity in the first preparatory dimension during delayed double-reach conditions with a 100 ms instructed pause. **e,** Activity in the first preparatory dimension during compound reach conditions. This panel plots the same data shown in Fig. 4c. **f,** Occupancy of all 20 preparatory dimensions during single reach conditions. This panel plots the same data shown in Fig. 4d. **g,** Occupancy of preparatory dimensions during delayed double-reach conditions with a 600 ms instructed pause. This panel plots the same data shown in Fig. 4e. **h,** Occupancy of preparatory dimensions during delayed double-reach conditions with a 300 ms instructed pause. **i,** Occupancy of preparatory dimensions during delayed-double reaches with a 100 ms instructed pause. **j,** Occupancy of preparatory dimensions during compound reach conditions. This panel plots the same data shown in Fig. 4f.

**Extended Data Fig. 7.**
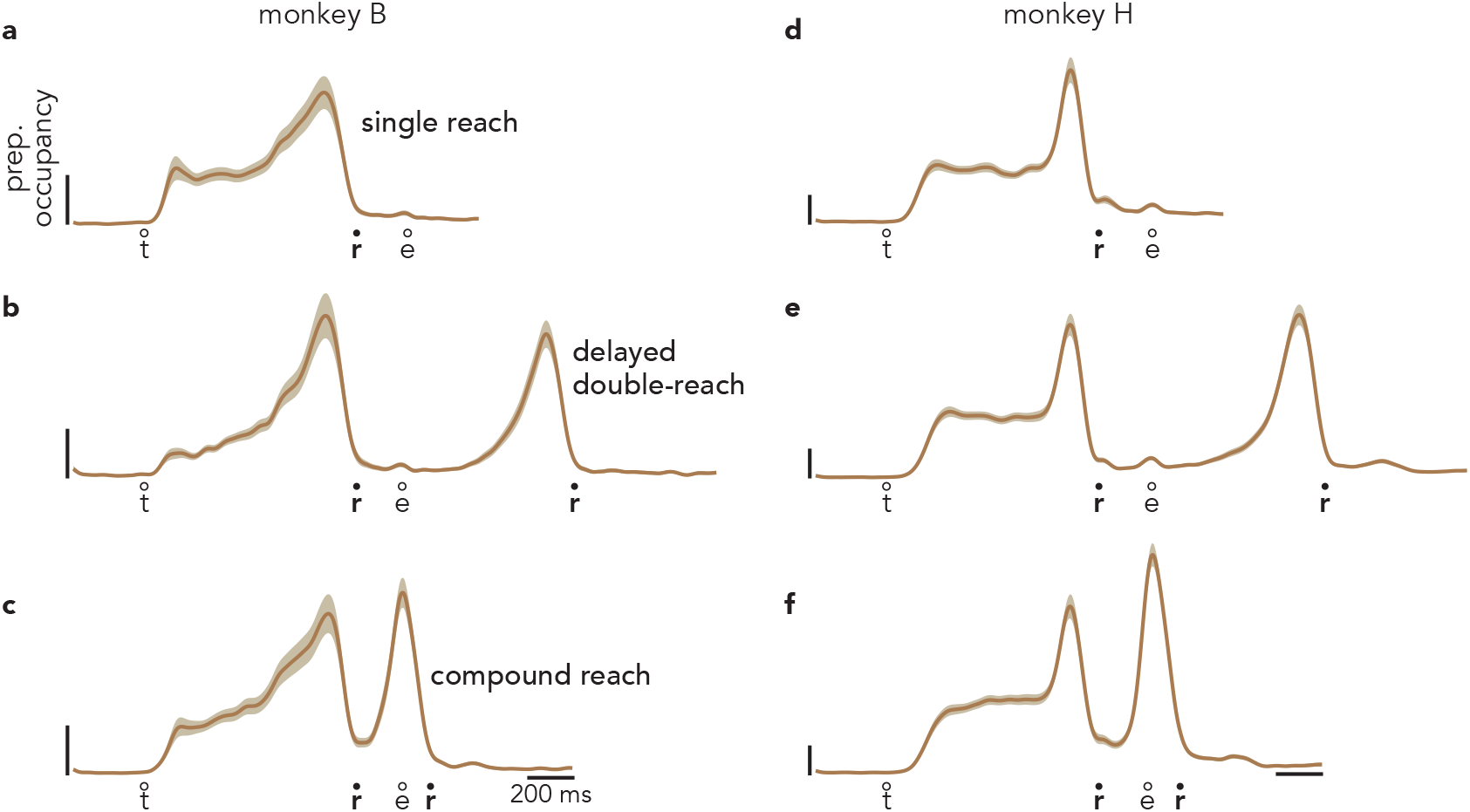
Defining the preparatory dimensions using all preparatory epochs. **a,** Similar to Fig. 4d, this panel plots the occupancy of 20 preparatory dimensions during single reach conditions. However, these dimensions were found using dwell-period activity from compound reach conditions (activity within a 40 ms window beginning 140 ms before the onset of the second reach) in addition to activity used to define the preparatory dimensions in Fig. 4d-f. Data shown are from monkey B. **b,** Occupancy of the same preparatory dimensions during delayed double-reach conditions. **c,** Occupancy of the same preparatory dimensions during compound reach conditions. This panel plots the same data shown in Fig. 4g. **d-f,** Same as **a-c,** but data are from monkey H.

**Extended Data Fig. 8.**
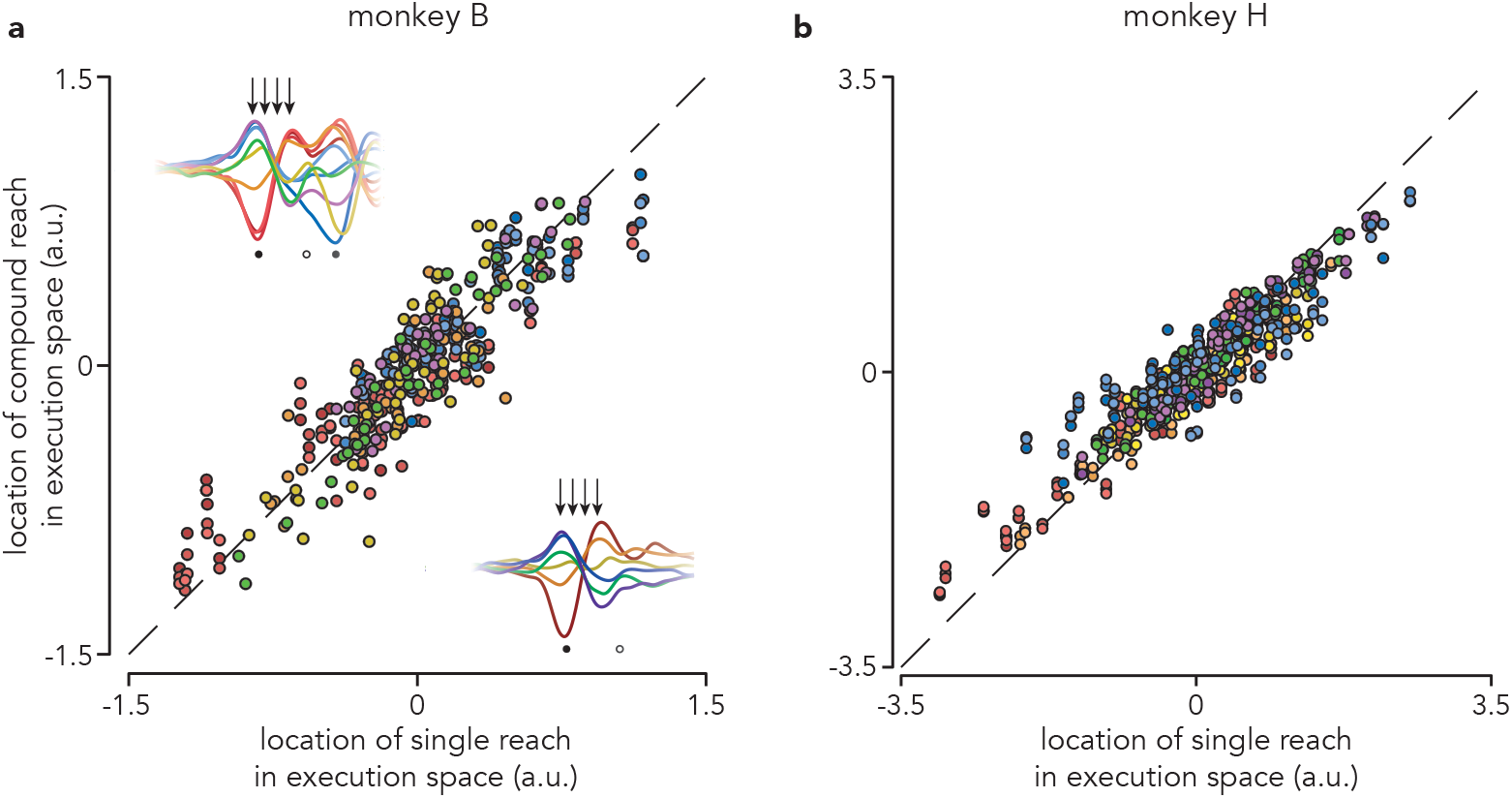
Patterns of execution-related activity. **a,** Comparison of activity within the top 10 execution dimensions during single and compound reach conditions. Similar to Fig. 5c, each marker represents the activity (in 1 of 10 execution dimensions, at a single point in time) of a pair of conditions that share a first reach. Unlike Fig. 5, here, we plot activity from 4 time points (50 ms intervals, starting 25 ms before reach onset). Data shown are from monkey B. **b,** same as **a**, but data are from monkey H. Although activity from the first reach of compound reaches was not used to define the preparatory and execution dimensions, these dimensions explained comparable amounts of variance during the first reach epoch of both single reaches and compound reaches. These dimensions explained 87% and 78% as much of the first reach epoch variance during compound reaches as was explained during single reaches, monkey B and monkey H, respectively.

**Extended Data Fig. 9.**
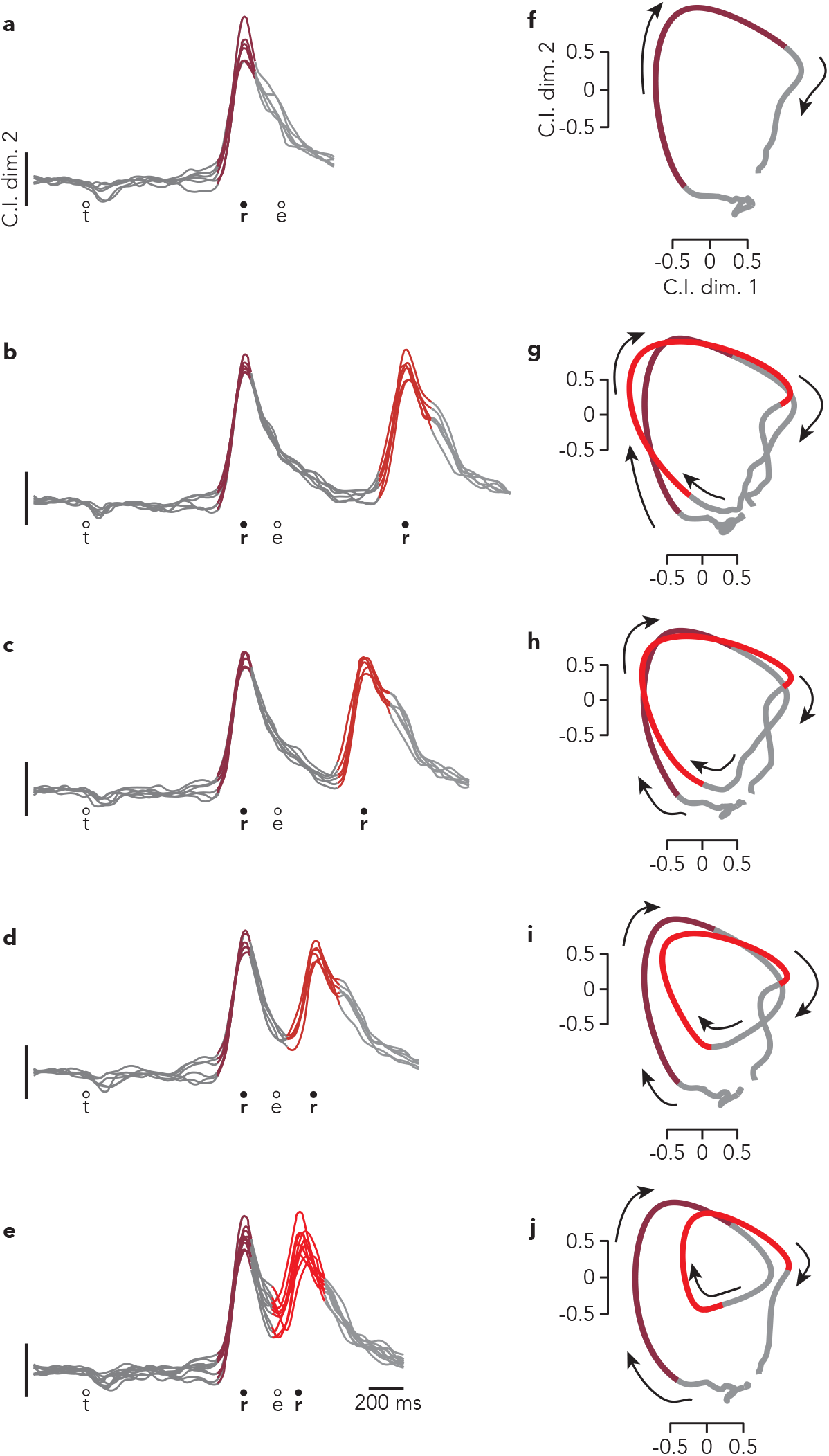
Activity within condition-invariant dimensions during all conditions, monkey B. **a,** Projections of population activity from single reach conditions onto a single condition-invariant dimension. As in Fig. 6, traces are colored to highlight peri-reach activity. This panel plots the same data shown in Fig. 6a, top. **b,** Same as a but for delayed double-reaches with a 600 ms instructed pause. This panel plots the same data shown in Fig. 6b, top. **c-d,** Same as **b** but for delayed double-reaches with 300 ms (**c**) and 100 ms (**d**) instructed delays. **e,** Same as **a** but for compound reaches. This panel plots the same data shown in Fig. 6c, top. **f,** Average (across-condition) activity in two condition-invariant dimensions during single reach conditions. This panel plots the same data shown in Fig. 6a, bottom. **g,** Same as **f** but for delayed double-reaches with a 600 ms instructed pause. This panel plots the same data shown in figure 6b, bottom. **h-i,** Same as **g** but for delayed double-reaches with 300 ms (**h**) and 100 ms (**i**) instructed delays. **j,** Same as **f** but for compound reaches. This panel plots the same data shown in Fig. 6c, bottom.

